# Macrophage-derived IL-27 sustains autoreactive CD8^+^ cytotoxic T cells in autoimmune hepatitis

**DOI:** 10.64898/2026.09.25.754422

**Authors:** Kaiyuan Hao, Sofia Grace Baptista, Aurora Stephania Virginia Beaumond, Daniel Milo Schafer, Kevin MingJie Gao, Prasanthi Lalitha Kaja, Kristy Chiang, Kerstin Nündel, Sushmitha Vijaya Kumar, Marc Samuel Sherman, Ann Marshak-Rothstein, Siva Karthik Varanasi

## Abstract

Chronic or persistent T cell activation is widely thought to culminate in T cell exhaustion, yet how autoreactive CD8⁺ T cells remain functionally intact despite sustained self-antigen exposure is poorly understood. In chronic infection and cancer this durability is attributed to stem-like, TCF1⁺ PD-1⁺ progenitor-exhausted (Tpex) cells that reside in lymphoid tissue and continuously replenish functional effectors; whether such a reservoir operates within a chronically inflamed peripheral organ during sterile autoimmunity is unknown. Here we show that mice lacking the lysosomal nuclease DNase2a together with the type I interferon receptor Ifnar1 (*Dnase2a⁻/⁻Ifnar1⁻/⁻*) develop spontaneous, progressive hepatic inflammation. This hepatitis was dependent on the endosomal DNA sensor TLR9, identifying self-DNA as the initiating ligand. Inflammation was marked by expansion of inflammatory macrophages and accumulation of CD8⁺ T cells co-expressing PD-1 and TOX that retained, rather than lost, effector function. Paired single-cell TCR and transcriptomic analysis revealed progenitor-exhausted T cells within the liver itself that shared clonotypes with clonally expanded PD-1^+^ TOX^+^ T cells, defining a locally operating progenitor-to-effector pipeline. Spatial transcriptomics positioned these intrahepatic T cell clusters adjacent to IL-27–expressing inflammatory macrophages, and disruption of IL-27 or its receptor in bone marrow–derived cells reduced PD-1⁺ CD8⁺ T cells and their effector functions. These findings define a novel mechanism for sustaining autoreactive T cells in the liver via TLR9-driven macrophage-induced IL-27 circuit that sustains a functional, self-renewing autoreactive PD1^+^ CD8⁺ T cell program in situ.

## INTRODUCTION

Autoimmune hepatitis (AIH) is a chronic, immune-mediated liver disease characterized by interface hepatitis, circulating autoantibodies, hypergammaglobulinemia, plasma cell clusters and, if untreated, progression to fibrosis, cirrhosis, and liver failure.^1,2^ Its hallmark is a persistent T cell-rich lymphoblastic infiltrate that sustains hepatocyte injury over years, and management still relies on non-specific corticosteroids and immunosuppression.^3,4^ Frequent relapse upon withdrawal underscores how incompletely the drivers of chronic intrahepatic T cell activation are understood.^5^ Progress has been constrained by a scarcity of tractable models: concanavalin A-induced and hepatocyte-antigen–induced hepatitis lead to acute, self-limited injury, or the death of the mice, and fail to recapitulate the chronic, spontaneous, progressive inflammation that defines human AIH.^6–8^ A model in which self-antigen drives durable, spontaneous hepatitis would allow the cellular circuits that maintain chronic hepatic autoimmunity to be dissected directly.

Endogenous nucleic acids are increasingly recognized as drivers of sterile autoinflammation, and their disposal is a core function of phagocytic macrophages.^9^ Genomic DNA released by the enucleation of erythroid precursors or by dying cells is engulfed by macrophages and normally degraded in phagolysosomes by the endonuclease DNase II; when this clearance fails, undegraded self-DNA accumulates within the very cells normally responsible for its removal and uncleared debris is then further internalized by additional proinflammatory antigen presenting cells.^9,10^ Loss of DNase II is embryonic lethal through an unrestrained type I interferon (IFN) response, and deletion of the type I IFN receptor (Ifnar1) rescues viability to yield *Dnase2a⁻/⁻ Ifnar1⁻/⁻* mice that are best known as a model of late onset inflammatory arthritis.^11,12^

Humans with biallelic DNASE2 mutations present with a systemic interferonopathy that includes hepatosplenomegaly and liver fibrosis.^13^ Self-DNA can be sensed through the cytosolic cGAS–STING pathway or through endosomal Toll-like receptor 9 (TLR9),^14^ and because inflammatory monocyte lineage cells both engulf self-DNA and express the endosomal DNA sensor TLR9,^15,16^ they are positioned to convert a clearance defect into a sustained inflammatory signal. Whether chronic self-DNA accrual causes AIH-like hepatitis, and through which sensor, has not been defined.

Persistent antigenic stimulation is classically thought to drive CD8⁺ T cells into exhaustion, a program instructed by the transcription factor TOX and marked by sustained PD-1 expression with progressive loss of effector function.^17,18^ Nevertheless, in autoimmune diseases including AIH, autoreactive T cells remain destructive across years of continuous self-antigen exposure.^19–21^ In chronic viral infection and cancer, durable effector responses are maintained by stem-like TCF1⁺ PD-1⁺ progenitor-exhausted (Tpex) cells that self-renew and replenish effectors from within lymphoid tissue.^22–26^ Whether an analogous progenitor sustains functional autoreactive CD8⁺ T cells in the chronically inflamed liver and which cells supply the factors that maintain these CD8 T cells remains unknown. Macrophages are strong candidates: beyond initiating inflammation, they serve as antigen-presenting cells and as a source of T cell-instructive cytokines in the liver.^27–29^

Here we introduce *Dnase2a⁻/⁻ Ifnar1⁻/⁻* mice as a spontaneous model of chronic autoimmune hepatitis. Disease strictly depends on the DNA sensor TLR9, and not TLR7, identifying self-DNA as the initiating ligand, and requires TLR9-expressing bone marrow-derived cells. The inflamed liver accumulates inflammatory monocyte-derived macrophages and hyperinflammatory CD8⁺ T cells co-expressing PD-1 and TOX that remain functionally potent rather than terminally exhausted. Paired single-cell TCR and gene-expression analysis identifies progenitor-exhausted T cells within the liver that share clonotypes with the clonally expanded effector pool, defining a locally operating progenitor-to-effector pipeline. Spatial transcriptomics localizes these T cell clusters adjacent to inflammatory macrophages expressing IL-27, and disruption of IL-27 or its receptor in bone marrow–derived cells reduce the number of PD-1⁺ CD8⁺ T cells and blunts their effector function. Together, these findings establish a chronic AIH model, identify TLR9 sensing of self-DNA as its trigger, and reveal a macrophage–IL-27 circuit that sustains a functional, self-renewing autoreactive CD8⁺ T cell program in the liver.

## RESULTS

### DKO mice spontaneously develop features of autoimmune hepatitis through a TLR9-dependent mechanism

Humans with loss-of-function mutations in DNASE2 develop liver disease associated with perivascular inflammation and fibrosis, features shared with autoimmune hepatitis (AIH). To determine whether *Dnase2⁻/⁻ Ifnar⁻/⁻* double-knockout (DKO) mice reproduce clinical features of human autoimmune hepatitis, we assessed liver pathology in detail. DKO mice spontaneously developed early-onset mononuclear infiltration surrounding intrahepatic vessels, indicative of interface hepatitis (Fig. 1A). By immunofluorescence, these infiltrates localized predominantly to the periportal region and in some cases near the central vein, whereas heterozygous (Het) littermate controls showed minimal infiltration (Fig. S1A). Immune infiltration was accompanied by extensive cell death (TUNEL⁺), age-progressive periportal fibrosis as detected by Sirius Red staining, elevated hepatic *Tnfa, Ifng*, *Cxcl10* by qPCR and increased serum ALT (Fig. 1B-F, S1B,C). In addition, DKO mice were hypergammaglobulinemic (Fig. 1G) and produced anti-nuclear autoantibodies (ANA) that stained HEp-2 cells with ANA patterns typical of Type I AIH (Fig. S1D,E). Thus, the DKO mouse recapitulates the diagnostic hallmarks of Type I AIH — interface hepatitis, bridging fibrosis, elevated ALT, and positive ANA.

**Figure 1.**
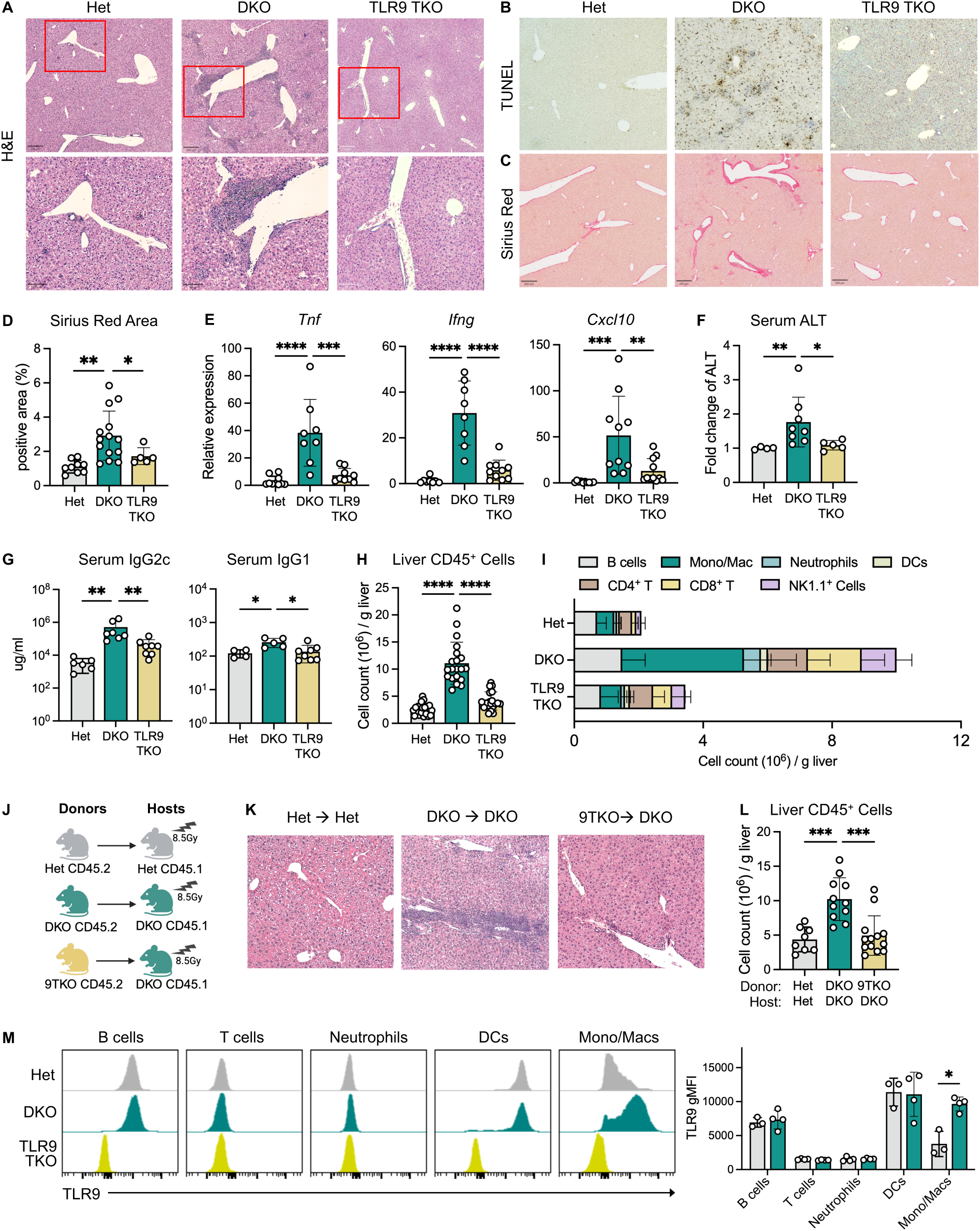
DKO mice spontaneously develop features of autoimmune hepatitis through a TLR9-dependent mechanism. (A) Representative H&E staining (scale bars, top panel 200um; bottom panel 100um) of liver sections from 3-4-month-old mice. (B) Representative TUNEL staining (scale bars, 200 um) of liver sections from 3-4-month-old mice. (C) Representative Sirius red staining (scale bars, 250 um) of liver sections from 3-4-month-old mice. (D) Quantification of Sirius red positive liver area ( n = 5-14 per group). (E) Relative hepatic expression of *Tnfa, Ifng, Cxcl10, and Col1a1* normalized to Het mice (n=8-10 per group). (F) Serum ALT level relative to Het controls in 3-4-month-old mice(n = 4-8 per group). (G) Serum total IgG1 and IgG2c levels in 3-4-month-old mice (n = 4-7 per group). (H) Absolute numbers of CD45^+^ immune cells in the livers of 3-4-month-old mice (n = 7-25 per group). (I) Absolute numbers of indicated immune population in the livers of 3-4-month-old mice (n = 9-14 per group). B cells (CD19+); CD4+ T cells (CD3+, CD4+); CD8+ T cells (CD3+, CD8+); NK1.1+ cells (NK1.1+, CD3-); DCs (CD11b-, CD11c+); Neutrophils (CD11b+, Ly6G+, Ly6C+); Monocytes/macrophages (CD11b+, Ly6G-, Ly6C+ / F4.80+). All cells were gated on live, CD45+, singlet cells. (J) Schematics of the bone marrow chimera experiment. Lethally irradiated (8.5Gy) CD45.1 recipient mice were reconstituted with 10^7^ bone marrow cells from CD45.2 donors as indicated. (K) Representative H&E staining (scale bars, 200um) of liver sections from chimeras 8-week post bone marrow transplant. (L) Absolute numbers of total CD45+ immune cells, in the livers of chimeras 8-week post transplant (n = 5-9 per group). (M) Histograms and quantification of TLR9 expression in indicated immune cell populations from the livers of Het and DKO, and TLR9 TKO mice. Data are representative of three independent experiments (n = 3–4 mice per group). Expression is reported as geometric mean fluorescence intensity (gMFI). Data are pooled from at least 3 independent experiments and represented as mean ± SEM. *p < 0.05, **p < 0.01, ***p < 0.001, and ****p < 0.0001

Prior work established that the cytosolic DNA sensor STING drives the embryonic lethality and deforming arthropathy of DKO mice through an NFκB-dependent pathway,^14,30^ whereas splenomegaly, cytopenia, elevated serum cytokines, autoantibody production, and liver fibrosis depend on Unc93B1,^31,32^ a chaperone protein that directs nucleic-acid– sensing TLRs to endolysosomal compartment. To confirm that STING was not required for liver disease and to identify the specific Unc93B1-dependent receptor responsible for liver inflammation, we intercrossed DKO mice with UNC93B-, STING-, TLR9- and TLR7-deficient strains. While STING-deficient DKO (DKO STING^gt/gt^) and TLR7-deficient DKO (TLR7 TKO) mice retained disease manifestations (Fig. S1F), interface hepatitis, fibrosis, cytokine production, elevated ALT were all UNC93B1-and TLR9-dependent (Fig. 1A–F, S1F). Additionally, TLR9 deficient mice had lower serum IgG and both TLR knockout strains failed to produce ANA (Fig. 1G, S1D).^31^ DKO mice develop splenomegaly due to extramedullary hematopoiesis. Spleen weights are significantly reduced in TLR9 TKO, but not TLR7 TKO mice (Fig. S1G). Together, these criteria identify TLR9 as the pivotal sensor in the development of hepatic inflammation in DKO mice.

### TLR9 expressing hematopoietic cells are required for the AIH phenotypes

We next defined the composition of liver immune infiltrates in Het, DKO and TLR9 TKO mice using spectral flow cytometry. The absolute number of total CD45⁺ immune cells dramatically increased in the DKO liver, with CD8⁺ T cells, NK1.1+ innate lymphocytes, and monocytes/macrophages showing the greatest proportional expansion relative to littermate Het controls (Fig. 1H, I). Importantly, TLR9-but not TLR7-deficiency significantly reduced the numbers of myeloid, NK1.1^+^ and CD8^+^ T cells, indicating the infiltration of immune cells to the liver was TLR9 dependent (Fig 1H, I, S1H).

To determine whether the TLR9-expressing cells that drive chronic inflammation were hematopoietic or stromal in origin, lethally irradiated DKO mice, were reconstituted with bone marrow stem cells from either DKO or TLR9 TKO donors (Fig. 1J), and the extent of AIH in these radiation chimeras was evaluated 8 weeks after reconstitution. DKO◊DKO positive-control chimeras recapitulated the DKO phenotype as confirmed by H&E and Trichome staining (Fig. 1K, S1I), establishing that the disease-promoting cell types were not lost during bone marrow transplantation. In contrast, TLR9 TKO◊DKO chimeras consistently failed to develop histological features of hepatitis with correspondingly reduced fibrosis and number of intrahepatic immune cells (Fig S1I, Fig 1L), particularly in the CD8⁺ T cell and myeloid compartments, relative to controls (Fig S1J). Thus, a TLR9-expressing bone marrow–derived cell is required for both hepatic and systemic inflammation of DKO mice.

To narrow the TLR9-expressing population that could contribute to the chronic liver inflammation in DKO livers, we compared TLR9 expression across liver immune cells by intracellular staining. While B cells, DCs, and monocytes/macrophages expressed TLR9, only minimal expression if not null, was observed in T cells and neutrophils (Fig. 1M). DNase II is required to process dsDNA-derived TLR9 ligands in B cells and DCs and therefore DKO B cells and DCs fail to respond to dsDNA under conditions where Het B cells and DCs mount vigorous responses^33,34^. Moreover, B cell and DC numbers were not proportionally expanded in the DKO liver (Fig. 1I) and expression levels of TLR9 in DKO and Het B cells and DCs were comparable. By contrast, TLR9 expression levels were much higher in DKO myeloid cells than in Het myeloid cells (Fig. 1M). Overall, these data suggest that TLR9 expressing monocytes/macrophages may drive liver inflammation during AIH.

### TLR9 signaling promotes inflammatory monocyte and macrophages accumulation during chronic liver inflammation

To characterize the phenotypic, functional, and transcriptional differences between the hepatic myeloid cells of Het and DKO mice, we performed single-cell RNA sequencing (scRNA-seq) on total liver immune cells (Fig 2A). Within the monocyte/macrophage compartment, six major subsets were resolved as defined by canonical marker genes described in previous reports (Fig 2B)^35–37:^ classical monocytes (Ly6c2, Ccr2, and Vcan), transitioning monocytes (*Slamf8, Slamf7,* and *Ldlr*), patrolling monocytes (*Pglyrp1* and *Ace*), monocyte-derived macrophages (MoMacs) (*Adgre1, Spn, Cd300e, Clec4b1,* and *Slamf9*), transitory macrophages (Arhgap22, Cdk14, Cadm1, and *Adgre1*), and Kupffer cells (Vsig4, Clec4f, and Timd4) (Fig. 2C). Whereas the myeloid compartment in Het livers was dominated by Kupffer cells, DKO livers were specifically enriched for all other inflammatory monocyte and macrophage subsets with MoMacs emerging as the most abundant myeloid population (Fig. 2B). Transitioning monocytes and MoMacs with comparable phenotypes have been described in other inflammatory liver conditions.^35–41^

**Figure 2.**
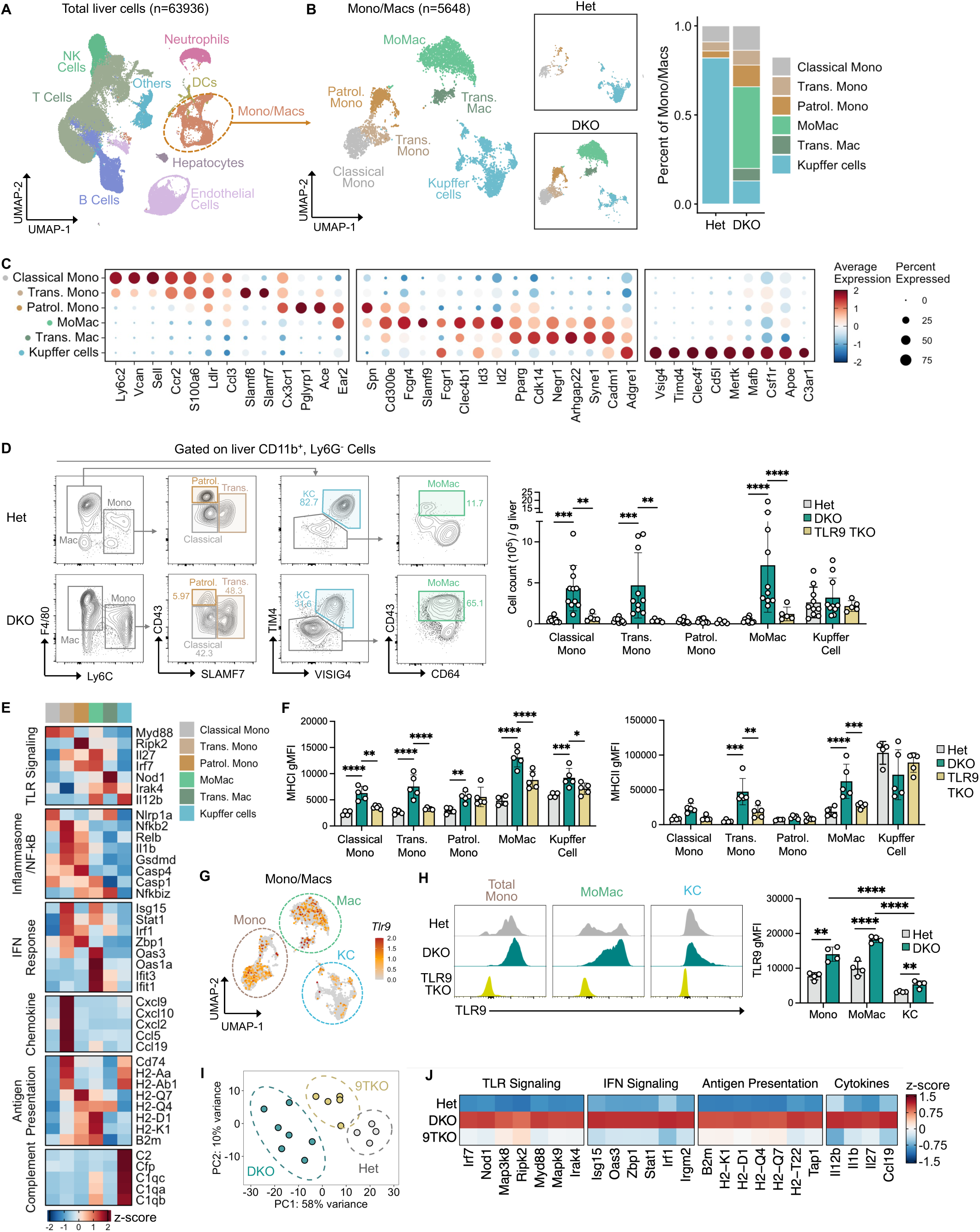
TLR9 signaling promotes inflammatory monocyte and macrophages accumulation during chronic liver inflammation. (A) Uniform manifold approximation and projection (UMAP) plots of total liver cells (n = 63,936 cells). (B) UMAP plots and relative abundance of monocyte/macrophage (Mono/Mac) subsets (n = 5,648 cells) in Het and DKO livers. Trans. Mono, transitioning monocyte; Patrol. Mono, patrolling monocyte; MoMac, monocyte-derived macrophage; Trans. Mac, transitory macrophage. (C) Expression of genes defining monocyte/macrophage subsets in (B). (D) Representative gating strategy to identify monocyte, macrophage subsets among liver CD3^−^ CD19^−^ Ly6G^−^ CD11b^+^ cells. Bar plot shows the absolute number of indicated populations in Het, DKO, and TLR9 TKO (n = 5-8 per group). (E) Heatmap showing RNA expression of indicated genes associated with TLR signaling, inflammasome and NF-κB signaling, IFN responses, antigen presentation, and complement across monocyte/macrophage subsets. Column colors denote classical monocytes (gray), transitional monocytes (beige), patrolling monocytes (brown), monocyte-derived macrophages (green), transitional macrophages (dark green), and Kupffer cells (blue), same as in Figure 2B. (F) Quantification of surface MHC I and MHC II expression in liver monocyte/macrophage subsets in Het, DKO, and TLR9 TKO livers (n = 4-5 per group). Expression is reported as gMFI. (G) Feature plot of Tlr9 RNA expression across monocyte/macrophage subsets. (H) Representative histograms and quantification of TLR9 expression in total monocytes, inflammatory macrophages, and Kupffer cells from Het, DKO, and TLR9 TKO livers (n = 4-9 per group). Expression is reported as gMFI. (I) Bulk RNA sequencing on sorted monocytes and macrophages (CD45+, CD3-, CD19-, CD11b+, Ly6G-, TIM4-) from Het, DKO, and TLR9 TKO livers (n=4-7 per genotype) was performed. Data shows Principle component analysis (PCA) of all samples. (J) Heatmap of DEGs (Het vs DKO) across all 3 genotypes. Data shown in (D), (F), and (H) are pooled from at least 3 independent experiments and presented as mean ± SEM. *p < 0.05, **p < 0.01, ***p < 0.001, and ****p < 0.0001. scRNA-seq data includes 2 liver samples per genotype.

Subset-specific surface markers as detected by flow cytometry confirmed the scRNA-seq data, resolving at least five myeloid subsets based on existing nomenclature: classical monocytes (LY6C^hi^), patrolling monocytes (LY6C^lo^, CD43⁺), transitioning monocytes (Ly6C⁺, SLAMF7⁺), MoMacs (LY6C⁻, CD64⁺, F4/80^+^, CD43⁺), and Kupffer cells (VSIG4⁺, TIM4⁺).^27^ Compared with Het controls, classical monocytes, transitioning monocytes, and MoMacs were particularly expanded by numbers and proportion in the livers of DKO mice, confirming the scRNA-seq findings (Figs. 2D, S2A). Importantly, the enrichment of monocytes and MoMacs was dampened but not completely reversed in TLR9 TKO mice (Figs. 2D, S2A). Together, these data identify a TLR9-dependent expansion of inflammatory monocytes and macrophages in the autoimmune liver environment.

Intriguingly, all myeloid subsets except for Kupffer cells expressed genes and pathways associated with TLR signaling (Irf7, Irak1, Nfkb1, and Il12), IFN responses (Isg15, Zbp1, Irf1, Oas1a), and inflammasome activation (Nlrp1a, Gsdmd, Casp1, and IL1b), indicative of their proinflammatory activation state (Fig. 2E). However, these myeloid subsets also exhibited distinct functional profiles as revealed by differentially expressed genes (DEG) and gene set enrichment analysis (GSEA). For example, transitioning monocytes preferentially expressed pathways and genes (*Cxcl9, Cxcl10, Ccl5*) associated with T/B cell chemotaxis that likely contribute to lymphocyte recruitment. Moreover, specific myeloid subsets upregulated genes associated with distinct antigen-presentation pathways: transitioning monocytes and Kupffer cells were enriched for MHC class II presentation (*Cd74, H2-Aa, H2-Ab1*), and patrolling monocytes and MoMacs for MHC class I (*H2-D1, H2-K1, B2m*) (Fig 2E, S2C). Flow cytometry resolved the same pattern showing that MoMacs in the DKO livers expressed higher levels of MHC I compared to the other subsets in a TLR9 dependent manner. Similarly, MHC II expression was upregulated in the transitory monocytes and MoMacs in the DKO livers also through TLR9-dependent mechanisms. However, Kupffer cells maintained high basal MHCII expression regardless of inflammation or TLR9 signaling (Figure 2F). Kupffer cells were enriched for genes and GO terms involved in complement activation (*C1qa, C2, Cfp*) and expressed numerous scavenger receptors, consistent with the key function of KC as clearance of complement coated bacteria from the intestine (Fig. 2E, S2B,D).^42^ Transitioning monocytes and MoMacs also expressed scavenger receptors involved in the clearance of dead cells (Fig. S2D). Overall, these findings demonstrate that DKO myeloid populations serve specific functions as far as antigen presentation capacity, cytokine and chemokine production, and clearance of cell debris.

We next assessed the contribution of TLR9 signaling to the transcriptional and functional changes in DKO myeloid cells. TLR9 expression, at both transcript and protein levels, was highest in inflammatory monocytes and macrophages and further induced by inflammation, whereas Kupffer cells expressed the least (Fig. 2G,H)^9^. We sorted monocyte-lineage cells (CD11b⁺Ly6G⁻TIM4⁻) from Het, DKO, and TLR9 TKO livers and performed bulk RNA-seq (Fig S2E). Principle component analysis revealed that TLR9-deficient cells clustered near Het controls (Fig. 2I) and loss of TLR9 led to downregulation of genes involved in inflammatory pathways characteristic of the DKO myeloid cells. These included genes involved in antigen presentation and as well as inflammatory mediators (*Il27, Il1b, Il12b*), critical for regulation of T cell responses (Fig. 2J). Thus, TLR9 signaling drives the recruitment and proinflammatory programming of monocytes and macrophages, which in turn may participate in recruitment and activation T cells in the liver during AIH.

### DKO livers harbor expanded PD-1+ TOX+ CD8+ T cells that exhibit effector function

Flow cytometry of the non-myeloid compartment showed that NK1.1+ cells and T cells, both major drivers of autoimmune liver disease and elevated in AIH patients,^43–47^ were also expanded in DKO livers through a TLR9-dependent mechanism (Fig 1I, 3A, Fig. S3A). The expanded NK1.1^+^ compartment in the DKO livers was composed mainly of CD49a⁺ DX5⁻ cells referred as Group 1 innate lymphoid cells (ILCs) or tissue-resident NK cells whereas Het livers contained mainly conventional NK cells (CD49a⁻ DX5⁺) (Fig S3B).^48–50^ Within the hepatic T cell compartment, CD8⁺ T cells were significantly enriched (8-9 fold) compared to CD4^+^ T cells (3-4 fold) (Fig 3A, S3A, S3C). CD8^+^ T cells from Het livers and spleens were predominantly naïve, while cells from DKO livers were almost entirely CD44^hi^ CD62L^lo^ activated cells (Fig. 3B, S3D). Within this CD44^hi^ population, approximately half of the cells expressed PD-1 and TOX with the rest comprising tissue-resident memory (T_RM) cells (CD69^+^, CXCR6^+^, and CD49a^+^) and short-lived effectors cells (T_SLEC) (KLRG1^+^) (Fig. 3B, S3E). Importantly, TLR9 TKO livers harbored a significantly reduced frequency of PD-1^+^ TOX^+^ CD8^+^ T cells, and an increased frequency of naïve CD8^+^ T cells, whereas T_RM and T_SLEC cell frequencies remained largely unchanged (Fig. 3B, S3F), indicating that TLR9-dependent signals are required for generating or maintaining PD-1^+^ TOX^+^ CD8^+^ T cells in the liver.

**Figure 3.**
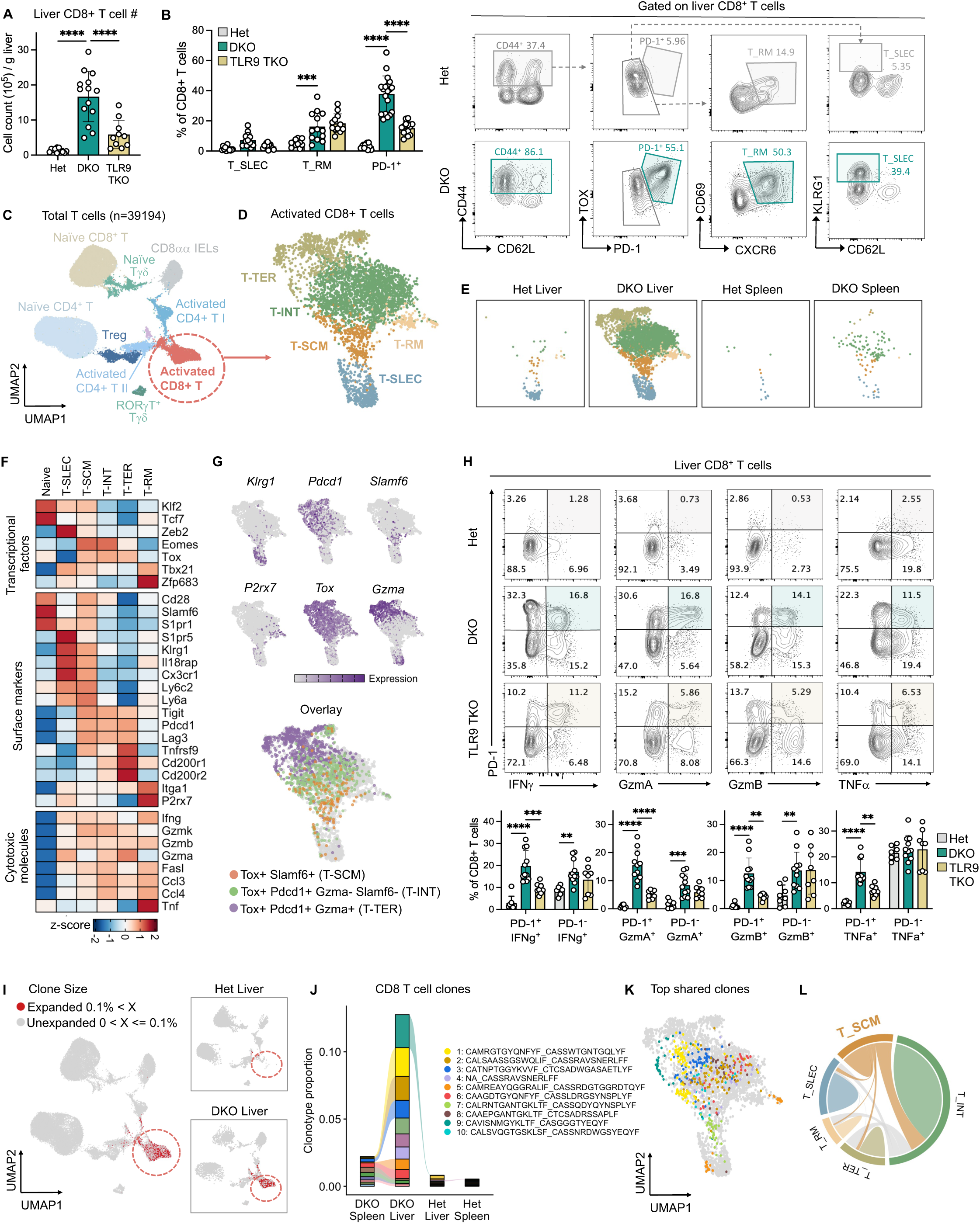
DKO livers harbor expanded PD-1+ TOX+ CD8+ T cells that retain effector function and undergo clonal expansion. (A) Absolute number of CD8^+^ T cells in livers of Het, DKO, and TLR9 TKO mice (n = 10-14 per group). (B) Representative gating strategy to identify PD-1^+^, tissue-resident memory (T_RM), and short-lived effector (T_SLEC) CD8^+^ T cell. Bar graph summarizes frequencies of each subset among liver CD8^+^ T cells (n = 11-19 per group). (C) UMAP plots of total T cells from the livers and spleens of Het and DKO mice (39,194 cells). (D) UMAP plots of activated CD8^+^ T cell (*Sell^neg^*) subsets from the livers and spleens of Het and DKO mice. (E) UMAP plots of activated CD8^+^ T cell subsets separated by tissue and genotype. (F) Heatmap showing RNA expression of transcription factors, surface markers, and cytotoxic molecules across different activated CD8+ T cell subsets. (G) RNA expression of key subset defining genes. Overlay shows T_SCM (*Tox, Slamf6*) and T_TER (*Tox,Pdcd1,Gzma*) cells. (H) Representative flow cytometry plots and frequencies of IFNγ, GzmA, GzmB, and TNFα-expressing PD-1^+^ and PD-1^−^ liver CD8^+^ T cells from Het, DKO, and TLR9 TKO mice post PMA and ionomycin stimulation. (I) UMAP plots of total T cells showing distribution of expanded TCR clones (clone size > 0.1%) and separated by genotype. (J) Top expanded CD8+ T cell clonotypes across tissues and genotypes, with each color representing one clone. (K) UMAP plot showing the distribution of the 10 most expanded TCR clonotypes shared with T_SCM population. Clonotypes also present in Figure 3J are shown in matching colors. (L) Chord diagram showing shared TCR clonotypes among activated CD8+ T cell subsets. Data shown in (A), (B), and (H) are pooled from at least 3 independent experiments and are presented as mean ± SEM. **p < 0.01, ***p < 0.001, and ****p < 0.0001. scRNA/TCR-seq data includes 2 liver and 1 spleen (pooled from two mice) per genotype.

To characterize the T cells further, we sorted on T cells (CD45^+^ CD3^+^) from liver and spleen of DKO and Het mice and performed paired single-cell gene expression and TCR sequencing. Unsupervised clustering resolved T cell clusters spanning αβ (CD8⁺, CD4⁺), γδ, and innate-like intraepithelial T cells (*Cd8a*⁺*Cd8b1*⁻*Cxcr6*⁺*Itga1*⁺) (Fig. 3C, S3G). Within CD4^+^ T cells, the proportion of Treg alone was significantly elevated in the DKO livers in a TLR9 dependent manner, whereas the other CD4⁺ T cell subsets such as Tbet^+^ Th1 and Rorgt+ Th17, and γδ^+^ T cell subsets remained largely unchanged as enumerated by scRNA and flow analysis (Fig. S3H,I). Interestingly, although composed of only a minor fraction of T cells, the number of CD8αα+ T cells that expressed tissue-residency markers (*Cxcr6* and *Itga1*) and *Gzmb* was elevated (although variable across mice) in the DKO liver, independently of TLR9 (Fig. S3J).

Sub-clustering the dominant CD8⁺ population identified five subsets: short-lived effectors (T_SLEC; *Klrg1*, *Cx3cr1*, *Zeb2*), resident memory (T_RM; *Itga1*, *P2rx7*, *Zfp683*), and three TOX⁺ PD-1⁺ subsets — stem-like memory (*Tcf1, Lef1, Slamf6*) (T_SCM), intermediate (*Lag3, Tigit, Eomes*) (T_INT), and terminally differentiated (*Cd200r1, Cd200r2, Tnfrsf9*) (T_TER) (Fig. 3D-F). Trajectory analysis supported a continuum from stem-like through intermediate to terminal states (Fig. S3K). This progression is marked by loss of SLAMF6 and gain of GzmA expression at both the protein and mRNA levels (Fig. 3G, S3L), resembling exhausted T cell differentiation in chronic viral infection and _cancer._23,51,52

We next assessed effector functions of these T cells. By gene expression, T_TER had the highest *Gzma*; T_INT were polyfunctional (*Gzmk*, *Gzmb*, *Gzma*, *Ifng*, *Fasl*, *Ccl3*, *Ccl4*); and T_SCM expressed *Gzmk*, *Gzmb*, and *Ifng*. (Fig. 3D). Intracellular staining confirmed polyfunctionality at the protein level as PD-1^+^ CD8^+^ T cells produced IFNγ and GzmA, and to a lesser extent GzmB and TNFα (Fig. 3H). IFNγ production by DKO liver CD8+ T cells was further validated by incorporation of an IFNγ reporter (GREAT mice) (Fig. S3M). Importantly, while the proportion of the effector cytokine (IFN-γ, GZMA, GZMB and TNF) producing PD-1+ CD8+ T cells were significantly reduced in TLR9 TKO livers, the cytokine producing PD-1-CD8+ T cells were unaffected by TLR9 deficiency, correlating with their ameliorated hepatitis (Fig. 3H). Together, these findings demonstrate that DKO liver promotes a TLR9-dependent expansion and differentiation of PD-1⁺TOX⁺ CD8⁺ T cells that exhibit robust effector functions.

### Development of AIH in DKO mice is IFNγ-dependent

Since IFNγ was one of the dominant effector cytokines produced by the multiple CD8⁺ T cell and other T and NK cell subsets in DKO livers (Fig S3G, S3N), it was of interest to determine how IFN-γ sensing contributed to DKO liver inflammation. We generated IFNγR TKO mice and found that these mice were completely protected from hepatitis, and showed no evidence of liver immune infiltration, fibrosis, or elevated serum ALT (Fig. S4A-C). This protection spanned all major expanded compartments: CD8⁺ T cells, NK cells, and monocytes/macrophages were all reduced relative to DKO controls (Fig. S4D-G). Moreover, the myeloid, NK, and T cell compartments resembled those of Het livers in composition, indicating that IFN-γ signaling contributes to the inflammatory compositional shift rather than merely for cell accumulation. Together, our data established IFNγ as a critical driver of sustained chronic hepatitis and liver damage, highlighting the importance of cytokine-producing cells in the DKO liver disease progression.

### Autoreactive cytotoxic T cells undergo clonal expansion in the DKO liver

To determine if the expansion of PD-1^+^ CD8^+^ T cells in the DKO liver reflected an antigen-driven response, we analyzed paired scTCR/RNA-seq data. This analysis identified extensive clonal expansion that was almost entirely restricted to activated CD8⁺ T cells, implicating antigen-specific responses in their chronic activation (Fig. 3I). Clonal expansion in CD4^+^ T cell compartment was comparatively minor (Fig. S5A). Of total CD8⁺ T cells, the 10 most expanded clones contributed to more than 10% of the repertoire; several expanded clones also appeared at low frequency in DKO spleen, suggesting inter-organ recirculation. However, no clonal expansion of T cells was observed in livers of Het littermates (Fig 3J). Within activated CD8^+^ T subsets, expanded clonotypes (∼50% of activated cells) spanned the T_SCM–T_INT–T_TER differentiation continuum, with the highly expanded clones (>1%) localized mainly to the T_TER cluster (Fig. S3B). These highly expanded CD8^+^ T cell clones were associated with expression of granzymes, most notably *Gzma*, and chemokines *Ccl3* and *Ccl4*, whereas *Ifng* expression was comparable across all activated CD8^+^ T cell subsets regardless of clone-size (Fig. S3C). This pattern was consistent with flow cytometry data showing IFN-γ production by both PD-1⁺ and PD-1⁻ CD8⁺ T cells, while GzmA production was preferentially associated with the PD-1⁺ population (Fig. 3H). Of all the expanded clones in the DKO livers, about 60% of clones were shared across different CD8^+^ T cell subsets. Observation of clones that can be found in more than one subset of CD8^+^ T cells further supported their differentiation continuum (Fig. S3D). Importantly, approximately 40% of stem-like CD8⁺ clones were shared with the T_INT population (Fig. 3K, L, S5E), supporting a role of stem-like T cells as precursors that sustain the clonally expanded cytotoxic T cells populations. Interestingly, these shared clones displayed substantial heterogeneity in effector programs, as individual clones expressed unique pattern of cytotoxic molecules such as *Gzma, Gzmb, Gzmk, Ifng, Ccl3, Ccl4* (Fig. S5F), suggesting that each clone could have a unique function and/or differentiated uniquely. Collectively, these data suggest that DKO livers harbor chronically activated, autoreactive CD8⁺ T cell clones that may arise from a progenitor pool in response to putative liver self-antigens.

### Macrophages are localized with CD8^+^ T cells within the liver during chronic inflammation

To identify the signals required for generating or maintaining clonally expanded CD8⁺ T cells in the liver during AIH, we profiled Het and DKO livers using Xenium Prime 5K spatial transcriptomics (5,000 + 50 additional genes) and transferred scRNA-seq labels onto the Xenium data. Using lineage-specific identity genes, we identified all major liver cell types along with their predicted locations, including hepatocytes, cholangiocytes, hepatic stellate cells, portal fibroblasts, mural cells, mesothelial cells, endothelial cells, and immune cells (macrophages, DCs, and T/B/NK cells) (Fig. 4A, S6A,B). We could also pick up clusters of plasma cells with abundant immunoglobulin kappa light chain transcripts (*Igkc*) (Fig. S6C). Comparing cellular abundance of non-hepatocytes (as hepatocytes were major proportion of cells in the liver) between Het and DKO livers revealed that DKO livers were particularly enriched in macrophage, DCs, and CD8^+^ T cells, confirming our scRNA-seq and flow cytometry findings (Fig. S6A).

**Figure 4.**
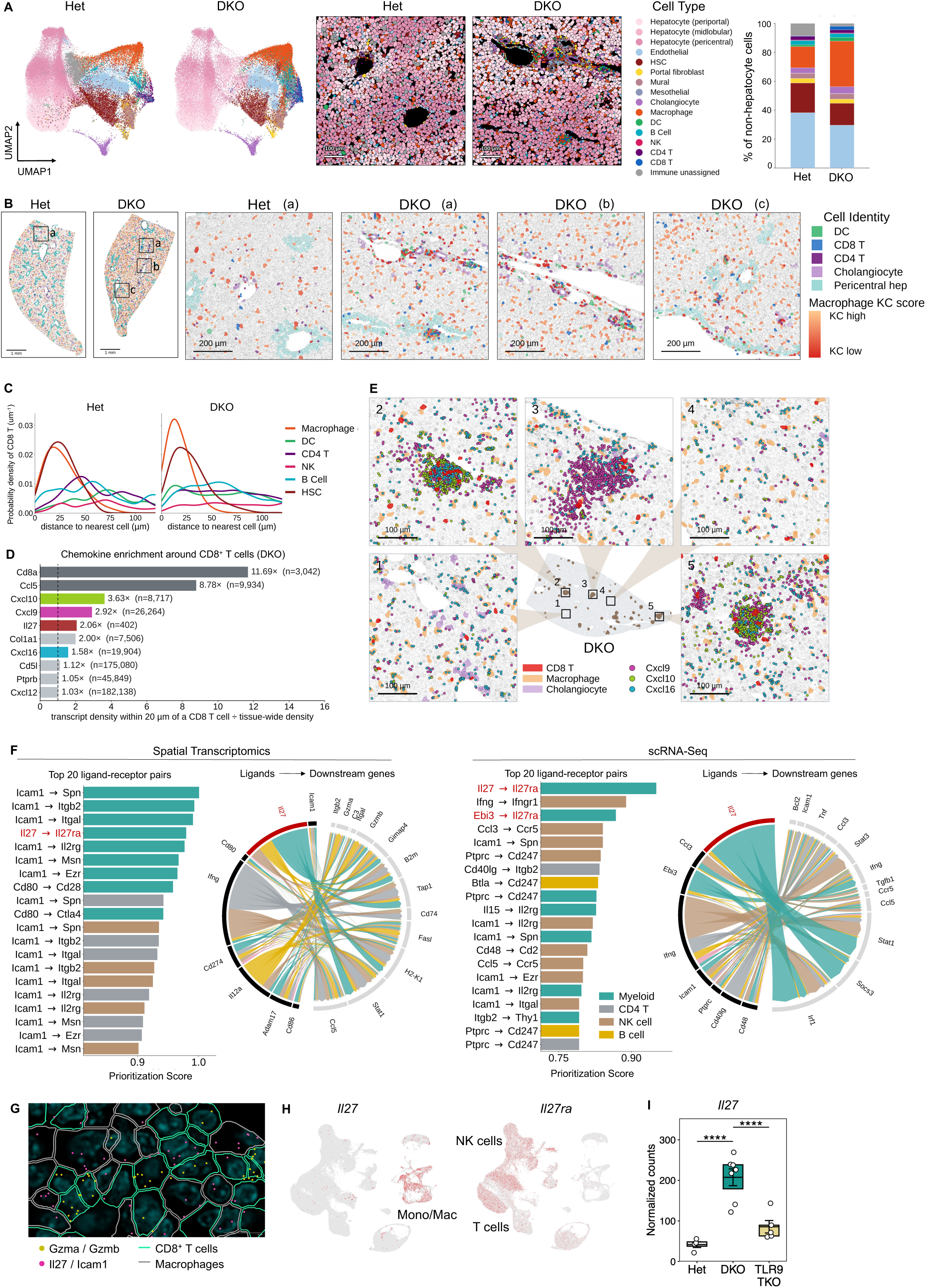
Inflammatory Macrophages are localized with CD8^+^ T cells within the liver in autoimmune hepatitis. (A) UMAP and representative spatial mapping of cells captured by Xenium spatial transcriptomics in Het and DKO liver sections. Staggered bar plot shows relative abundance of non-hepatocytes cells. (B) Xenium spatial transcriptomic analysis showing the distribution of the indicated immune-cell populations in liver sections from Het and DKO mice. Boxed regions in the tissue overview (left) (scale bar, 1mm) are shown at higher magnification (right) (scale bar, 200um). Macrophages are colored according to their continuous Kupffer cell (KC) score. Cholangiocytes and pericentral hepatocytes delineate portal and central regions, respectively. (C) Distribution of nearest-neighbor distance between CD8⁺ T cells and the indicated cell populations in Het and DKO liver sections. Curves represent Gaussian kernel-density estimates. (D) Fold enrichment of the indicated transcripts within 20 um of CD8+ T cells relative to their tissue-wide densities in DKO liver section. Dash line denotes cut-off for no enrichment (fold change = 1). n indicates the total number of detected transcripts for each gene. (E) Spatial distribution of *Cxcl9*, *Cxcl10*, and *Cxcl16* transcripts and the indicated cell populations in the DKO liver section. The whole-tissue overview shows the locations of five zoomed regions. (F) NicheNet analysis of cell-cell interactions inferred from spatial transcriptomics and scRNAseq data. Bar graph shows the top 20 predicted ligand–receptor pairs targeting CD8+ T cells. (G) Representative spatial transcriptomic image showing macrophage and CD8^+^ T cell interactions in the DKO liver section (scale bar, 10um). (H) UMAP plots of total liver icells showing *Il27* and *Il27ra* expression as identified by scRNAseq. (I) Expression of Il27 in sorted monocytes and macrophages (CD45+, CD3-, CD19-, CD11b+, Ly6G-, TIM4-) from Het, DKO, and TLR9 TKO livers as determined by bulk RNAseq . ****p < 0.0001.

Hepatic functions are spatially segregated along the portal–central axis of the liver lobule, a phenomenon known as liver zonation. Accordingly, hepatocytes display transcriptional heterogeneity across this axis and can be classified as portal (near the portal vein), central (near the central vein), or mid-lobular (in between).^53–55^ We used *Glul* as a marker of central-vein hepatocytes and identified ∼10 genes that are most correlated and anticorrelated with *Glul*,^56^ defining them as pericentral and periportal landmark gene sets, respectively (Fig. S6D). The normalized summed expression of these sets was used to assign each hepatocyte a percentile-based zone index (Fig. S6E). While hepatocytes in Het livers were evenly distributed across portal, central, and mid-lobular zones, hepatocytes in DKO livers were predominantly periportal, indicating altered hepatic zonation during AIH (Fig. S6E,F).

Notably, inflammatory infiltrates characterized by accumulation of T cells, macrophages, and dendritic cells (DCs) were observed throughout the DKO liver, particularly around the portal and central veins (Fig. 4B). Within these infiltrates, nearest-neighbor analysis revealed that CD8⁺ T cells were more closely associated with macrophages than with DCs (Fig. 4C). Additionally, CD8⁺ T cells expressing *Gzma*, *Gzmb*, and *Gzmk* were observed within macrophage clusters (Fig. 4B, S6G).

Next, we determined which macrophage subsets were associated with CD8⁺ T cells. Because the Xenium 5K panel probes a limited gene set, we could not resolve distinct macrophage subsets directly as we did with scRNA seq. Instead, we scored macrophages (*Adgre1⁺, Aif1⁺*) using Kupffer cell (KC) identity genes (*Vsig4* and *Cd5l)* and classified macrophages as KC-high or KC-low based on this score (Fig. S6H). Our analysis indicated that KC-high macrophages were located predominantly in the midlobular region, whereas KC-low macrophages were found near the portal and central venules and in closer proximity to CD8⁺ T cells (Fig 4B). Consistent with this analysis, immunofluorescence imaging localized CD8⁺ T cells adjacent to hepatic macrophages, clustering specifically with CD4⁺ T cells and F4/80^lo^ macrophages rather than F4/80^high^ Kupffer cells (Fig. S6I).

To identify factors promoting CD8⁺ T cell accumulation in the DKO liver, we quantified transcript density within 20 µm of each CD8⁺ T cell, normalized to tissue-wide density. This analysis identified, in addition to T cell–associated genes (*Cd8a*, *Ccl5*), several chemokines (*Cxcl9*, *Cxcl10*, *Cxcl16*) and cytokines (*Il27*) specifically enriched in CD8⁺ T cell–rich regions (Fig 4D,E). Similarly, NicheNet analysis, performed independently on the spatial transcriptomic and scRNA-seq datasets, identified that monocyte/macrophage-derived signals including *Icam1*, *Il27*, and *Ebi3*, as leading candidates driving differential gene expression in CD8⁺ T cells from DKO livers (Fig. 4F). Furthermore, both *Il27 and Icam1* were expressed by macrophages in close contact with CD8⁺ T cells as shown by spatial transcriptomics (Fig. 4G). Notably, *Il27* was expressed exclusively by monocytes and macrophages and was upregulated in the MoMacs in the DKO liver, while *Il27ra* was expressed predominantly by T cells, as shown by scRNA-seq (Fig. 4H, S6J). Moreover, this upregulation of *Il27* in DKO liver myeloid cells was TLR9-dependent (Fig. 4I). Together, these data suggest that TLR9 dependent macrophage-derived IL-27 in the DKO liver interacts with CD8⁺ T cells to promote their differentiation.

### Macrophage-derived IL-27 sustains autoreactive CD8⁺ T cells and drives hepatitis

To determine how IL-27 signals within CD8⁺ T cells, in vitro–activated CD8⁺ T cells were exposed to IL-27, IL-15, or IL-2 (control). While both IL-27 and IL-15 induced an exhaustion program (*Pdcd1, Tox*), only IL-27 and not IL-15 stimulated cells displayed enhanced terminal differentiation genes (*Gzma, Tnfrsf9, Gzmb, Gzmk*) alongside stemness genes (*Tcf7, Slamf6, Ly6a*) (Fig. 5A).

**Figure 5.**
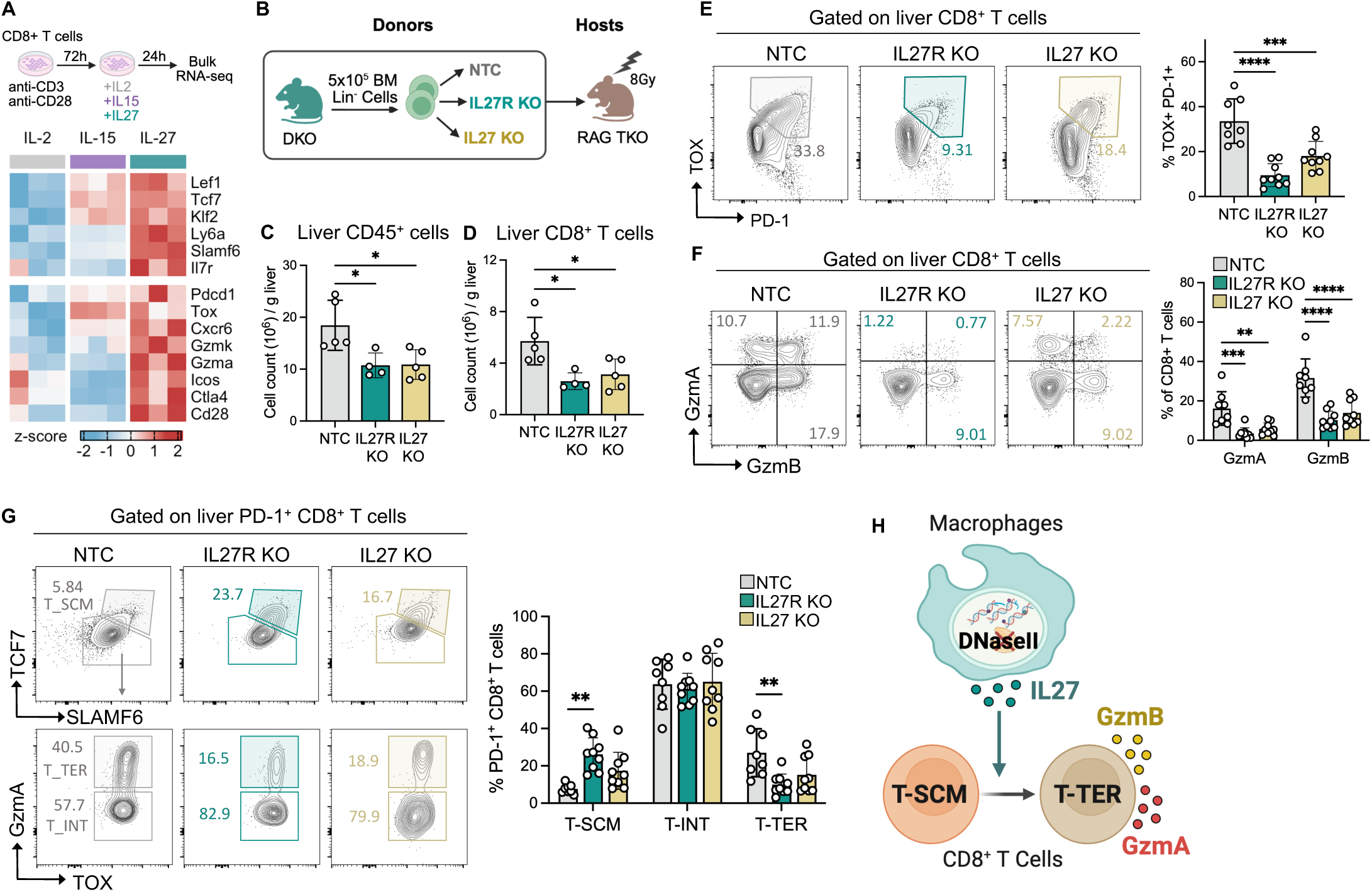
IL-27 sustains autoreactive CD8⁺ T cells and promotes T cell terminal differentiation. (A) Heatmap showing expression of genes associated with stemness (top) and differentiation (bottom) in *in vitro* cultured WT CD8^+^ T cells treated with indicated cytokines post TCR stimulation. (B) Schematics of the bone marrow chimera experiment. DKO Lin^-^ bone marrow cells were subjected to CRISPR-Cas9 mediated gene knockout as indicated and injected into lethally irradiated (8Gy) RAG TKO hosts. (C) Absolute numbers of total CD45^+^ immune cells in the livers of chimeras 9-week post transplant (n = 4-5 per group). (D) Absolute numbers of CD8^+^ T cells in the livers of chimeras 9-week post transplant (n = 4-5 per group). (E) Representative flow plots and frequencies of PD-1^+^ TOX^+^ cells among total liver CD8^+^ T cells from bone-marrow chimeras 9-week post transplant (n = 8-9 per group). (F) Representative flow plots and frequencies of GzmA and GzmB producing cells among total liver CD8^+^ T cells from bone-marrow chimeras 9-week post transplant (n = 8-9 per group). (G) Representative gating strategy and frequencies of PD-1^+^ CD8^+^ T cells subsets in the livers of bone-marrow chimeras 9-week post transplant (n = 8-9 per group). (H) Proposed model showing macrophage-derived IL27 promotes differentiation of T-SCM into T-TER cells and enhances granzyme production. Data shown in (C) and (D) are representative of 2 independent experiments; data shown in (E) - (G) are pooled from 2 independent experiments and presented as mean ± SEM. *p < 0.05, **p < 0.01, ***p < 0.001, and ****p < 0.0001.

We next assessed whether IL-27 signaling was necessary for CD8⁺ T cell responses in the liver during AIH. To this end, we reconstituted lethally irradiated RAG TKO hosts with DKO bone marrow stem cells in which *Il27* or *Il27ra* had been deleted by electroporation of a Cas9–gRNA ribonucleoprotein complex (Fig. 5B). At 8 weeks post-chimera, knockout efficiency was confirmed by IL-27R surface staining on circulating T cells (Fig. S6K) and changes in immune cell populations were enumerated. Deletion of either *Il27* or *Il27ra* in hematopoietic stem cells markedly reduced immune cell infiltration in the liver (Fig. 5C), particularly CD8⁺ T cells as reflected by decreased proliferation and absolute cell count (Fig. 5D, S6L). Importantly, knockout of *Il27* or *Il27ra* reduced the frequency of PD-1⁺TOX⁺ CD8⁺ T cells and their production of granzymes (GZMA/GZMB) (Fig. 5E,F). This blunted CD8^+^ T activity was independent of Treg abundances, as the Treg frequencies remained unchanged (Fig. S6M). Notably, *Il27* or *Il27ra* knockout CD8^+^ T cells displayed increased T_SCM (PD-1⁺TCF7⁺SLAMF6⁺) cells and reduced T_TER (PD-1⁺GZMA⁺TCF7⁻) cells (Fig. 5G), suggesting that macrophage-derived IL-27 signaling promotes terminal differentiation and of progenitor CD8⁺ T cells (Fig 5H). Together, these data establish that TLR9-driven, macrophage-derived IL-27 sustains the autoreactive CD8⁺ T cell response.

### AIH patients display PD-1+ TOX+ T cells with effector potential

To assess the relevance of this inflammatory circuit to human disease, we reanalyzed recently published single-nucleus RNA-seq (snRNA-seq) data from liver biopsies of patients with various liver diseases including AIH, PBC and MASH, and healthy controls.^57^ Within the CD8⁺ T cell compartment, a TOX⁺ CD8⁺ T cell population was uniquely expanded in AIH livers and was phenotypically similar to the PD-1⁺TOX⁺ CD8⁺ T cells enriched in DKO livers (Fig. 6A). These AIH TOX⁺ CD8⁺ T cells showed features of terminal differentiation (*TNFRSF9*, *DGKH*, *TIGIT*) and reduced expression of stemness-associated genes, yet retained residual effector potential (*IFNG*, *CCL4*) (Fig. 6B). A separate cluster with stem-like features (*SLAMF6*, *TCF7*, *LEF1*) and a stronger cytotoxic program may represent a precursor to the differentiated TOX⁺ population. Cross-species comparison revealed substantial transcriptional overlap between TOX⁺ CD8⁺ T cells in AIH livers and PD-1⁺ CD8⁺ T cells in DKO livers, supporting enrichment of PD-1⁺TOX⁺ CD8⁺ T cells in both human AIH and the DKO model (Fig. 6C). Consistent with these findings, analysis of peripheral blood mononuclear cells (PBMCs) from patients with AIH showed increased NK cells and cytotoxic CD8⁺ T cells (Fig. S7A,B) that expressed TOX PD-1, and SLAMF6, and together with multiple effector molecules, including *GZMA*, *GZMH*, *GZMM*, *GZMK*, *PRF1*, *CCL4*, *CCL5*, and *IFNG* (Fig. S7C),^58^ mirroring the PD-1⁺TOX⁺ effector program in DKO livers.

**Figure 6.**
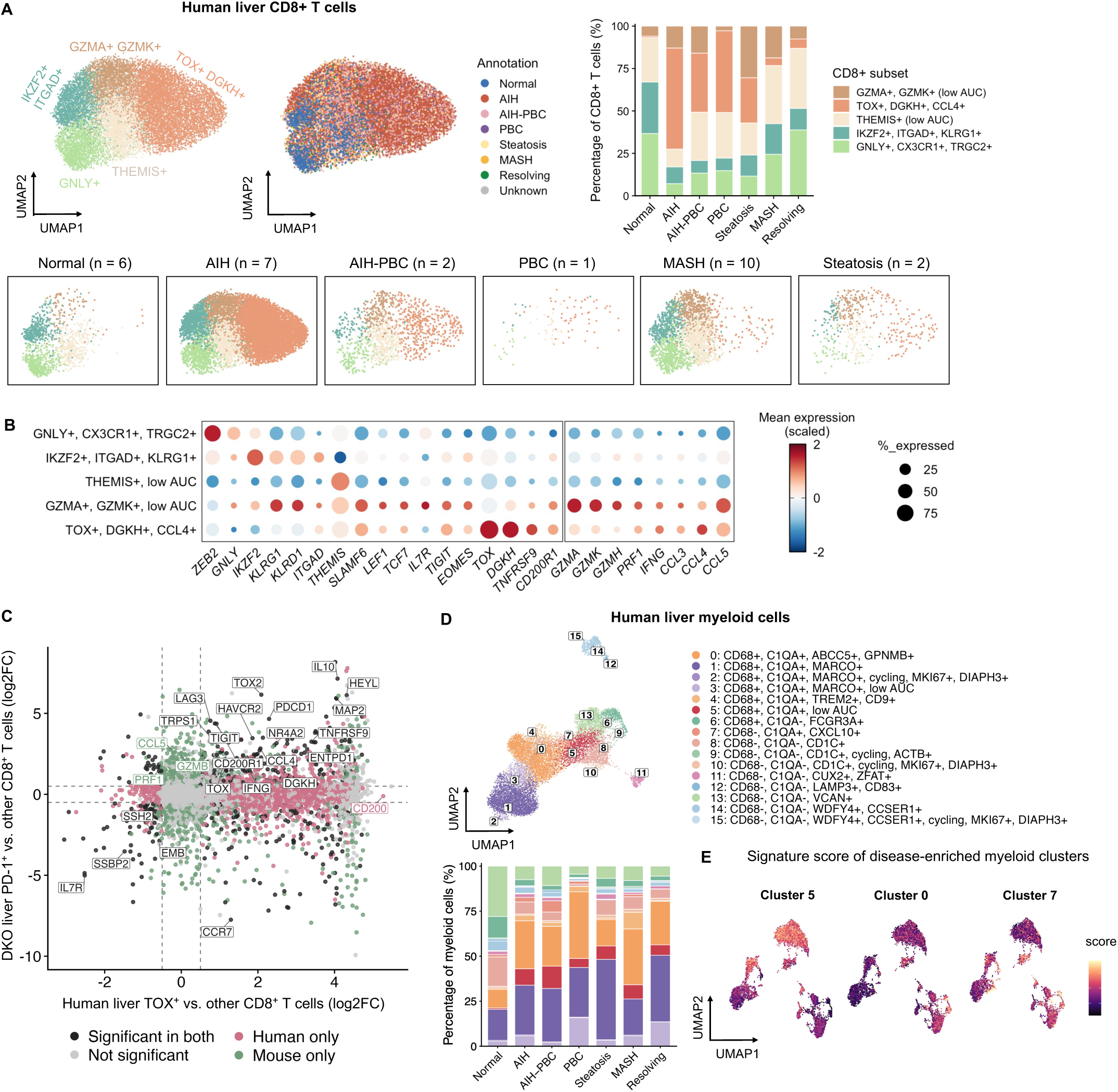
AIH patients display PD-1+ TOX+ T cells with effector potential. (A) UMAP plot of liver CD8+ T cell subsets and their distribution across the indicated diagnosis groups in single-nucleus RNA-seq dataset. Stacked bar graph shows the relative abundance of each CD8+ T cell subset by diagnosis. (B) RNA expression of cell-state-defining and effector genes across the CD8+ T cell subsets identified in (D). (C) Scatterplot compares differential gene expression in human liver TOX+ CD8+ T cells versus other human CD8+ T cell subsets with that in DKO liver PD-1+ CD8+ T cells (T_TER, T_INT, and T_SCM) versus the remaining DKO CD8+ T cells. Colors indicate genes significantly differentially expressed in both datasets (black), human only (pink), or mouse only (green). (D) UMAP plot of human liver myeloid cells annotated by the indicated clusters. Stacked bar graph shows the relative abundance of each myeloid-cell cluster across the indicated diagnoses. (E) UMAP projection of the gene module scores associated with the AIH-enriched myeloid-cell clusters. Cluster 5 module score comprises *Rtn1, Itga4, Arhgap22, Cst3, Txnip, H2-D1,* and *H2-Q7* (AUROC ≥ 0.65, %FC ≥ 1.5, log2FC > 0.8); cluster 0 module score comprises *Abcc5, Gpnmb, Fmn1, Slc1a3, Hs3st2, Cyp27a1, Myo1d,* and *Cpm (*AUROC ≥ 0.65, %FC ≥ 1.5, log2FC > 1); cluster 7 module score comprises *Nfkb1, Dram1, P2rx7, Sod2, Slamf7, Cd83, Arhgap31, Traf3, Pstpip2, Rnf144b, Acsl1, Rtn1, Wars1, Cd44, Icam1, Slc1a3, Hivep1, Cxcl10, Gch1, Rasgef1b, Niban1, Alcam, Zc3h12c, B4galt5,* and *Mgll* (AUROC ≥ 0.65, %FC ≥ 1.5, log2FC > 1).

We next asked whether DKO monocyte/macrophage populations share features with myeloid cells enriched in diseased human livers. We compared myeloid subsets identified by snRNA-seq across disease groups (Fig. 6D) and projected the differentially expressed gene signatures of each subset onto mouse monocyte/macrophage clusters (from Het and DKO). Relative to healthy livers, Cluster 5 (*CD68*⁺*C1QA*⁺, low AUC) and Cluster 0 (*CD68*⁺*C1QA*⁺*ABCC5*⁺*GPNMB*⁺) were proportionally expanded across liver diseases, whereas Cluster 7 (*CD68*⁺*C1QA*⁺*CXCL10*⁺) was preferentially expanded in AIH.^57^ Cluster 5 signatures were enriched in MoMacs, transitory macrophages, and patrolling monocytes, all populations expanded in DKO livers. Cluster 0 signatures were enriched in transitory macrophages and a subset of Kupffer cells; and Cluster 7 signatures were enriched in transitioning monocytes, consistent with their expression of T cell–recruiting chemokines (Fig. 6D,E). Together, these analyses indicate that the CD8⁺ T cell program and macrophage features are highly conserved between mouse and human AIH, and that the *Dnase2a*-deficient model recapitulates key features of the human disease.

## Discussion

Our study links inflammatory macrophages and their production of IL-27 to sustaining an effector autoreactive T cell response in the liver during autoimmune hepatitis. We introduce *Dnase2a⁻/⁻ Ifnar1⁻/⁻* (DKO) mice as a spontaneous, chronic model of autoimmune hepatitis (AIH) that reproduces the cardinal features of human disease, interface hepatitis, elevated ALT, progressive fibrosis, ANA, hypergammaglobulinemia, and hepatic plasma cell clusters and traces them to TLR9 sensing of self-DNA or microbial DNA by hematopoietic cells. Disease is driven by a self-reinforcing tissue circuit in which TLR9⁺ inflammatory macrophages produce IL-27 that promotes the differentiation of intrahepatic TCF1⁺ progenitors into CD8 effector cells, whose IFN-γ in turn amplifies macrophage activity. Because self-DNA release and TLR9 sensing also drive non-alcoholic steatohepatitis, alcoholic liver disease, and ischemia–reperfusion liver injury,^15,59–61^ this macrophage–IL-27–CD8⁺ T cell loop may represent a shared axis of chronic liver inflammation and a target that spares the global immunosuppression on which current AIH management depends.

How self-DNA is sensed to ignite this hepatic circuit, however, proved unexpectedly complex. Prior studies identified STING, a cytosolic DNA sensor, as responsible for the embryonic lethality of DNAse2a^-/-^ mince and the inflammatory arthritis that develops in DKO mice.^14,30^ We now confirm that endosomal nucleic acid sensors govern all aspects of DKO hepatitis; TLR7 was previously implicated in the production of ANAs,^31^ and we now document a critical role for TLR9 in all other manifestations of hepatitis. This outcome most likely reflects the dual role of DNase II – it both limits DNA accumulation in phagocytic cells while at the same time generating the short oligonucleotides that engage the TLR9 binding site in B cells and pDCs.^33,34,62,63^ The current study suggests that myeloid cells use additional nucleases to process DNA, so DNA-derived nucleotides are available to engage TLR9 independently of DNase II. The dependency of different tissues (joint and liver) and cell types (B cells and myeloid cells) on distinct nucleic acid sensors points to the importance of considering tissue or cell type specific therapeutic strategies even in monogenic diseases.

The ability to interrogate this self-DNA–TLR9–macrophage axis in a spontaneous setting distinguishes our model from prior systems. While many models of AIH have been proposed, ours recapitulates type 1 AIH more faithfully. The most commonly studied model, Con A-induced hepatitis is acute, is not triggered by autoantigens, and fails to progress to fibrosis.^6–8,64^ Pseudoautoantigen and liver-specific protein (CYP2D6/FTCD) immunization models rely on exogenous adjuvants and lead to CD4-driven diseases that recapitulate type II AIH.^65^ These models work best on NOD backgrounds that include numerous autoimmune susceptibility genes and are at odds with patient data identifying PD-1⁺TOX⁺ CD8⁺ T cells as a defining feature of AIH.^7,8,57,66^ The spontaneous PD-1⁺ TOX⁺ CD8⁺ response in DKO livers, arising from a defined trigger on an intact repertoire, addresses these gaps.

This faithful CD8⁺ T cell response, in turn, allowed us to ask where autoreactive PD-1⁺ TOX⁺ CD8⁺ T cells arise. While their origin and priming remain an open question, stem-like progenitors residing in the spleen and lymph nodes are thought to continuously give rise to differentiated effector cells. However, we find that intrahepatic CD8⁺ T cells contain stem-like clones that share TCR repertoires with the intermediate and terminally differentiated CD8⁺ T cells, arguing that the liver itself, rather than secondary lymphoid organs, may generate effectors from a local TCF1⁺ stem-like memory reservoir. Indeed, we identify IL-27 as the driver of this progenitor-to-effector transition: loss of IL-27R or IL-27 increased the frequency of TCF1⁺ cells while reducing granzyme A/B-expressing effectors in the liver, indicating that IL-27 either pushes stem-like cells to differentiate into PD-1⁺ TOX⁺ effectors or promotes the survival of the later rather than simply expanding the number of differentiated effector subsets as previously reported during chronic viral infection.^67^ In addition to IL-27, recent reports indicate that IL-15 and IL-21 can also promote the differentiation of progenitor-exhausted cells into effectors;^68–70^ however, we did not detect Il15 or Il21 near CD8⁺ T cell clusters by spatial transcriptomics, and our NicheNet analysis, which considered these and other candidate ligands that were both differentially upregulated and predicted to signal to CD8⁺ T cells, identified macrophage-derived IL-27 as the dominant niche signal. Importantly, IL-27 in chronic viral infection promotes stem-like cell expansion in the spleen,^67^ whereas in autoimmune liver disease it may instead promote terminal differentiation. This tissue-specific effect of IL-27 is further supported by a recent report that, during chronic hepatitis B virus (HBV) infection, CD4⁺ T cells license Kupffer cells to produce IL-27 that is essential for CD8⁺ T cell expansion and function;^29^ there, IL-27 blockade reduced HBV-specific T cell numbers and effector responses in the liver, while recombinant IL-27 increased the number of functional (GZMB⁺ IFN-γ⁺) HBV-specific T cells. Importantly, we find that the IL-27 producing cells in the DKO liver are predominantly inflammatory macrophages rather than Kupffer cells. Although the relative contribution of these two populations in CD8⁺ T cell differentiation in the liver will need to be resolved in future studies.

A pro-inflammatory function for IL-27 in this setting may at first appear to contradict its well-established immunoregulatory activities where IL-27 was previously shown to restrain Th2 and Th17 responses and promotes IL-10–producing Tr1 and induced regulatory T cells,^71–73^ functions that have framed it primarily as an anti-inflammatory cytokine. Yet IL-27 has also been reported to suppress Foxp3 induction in vitro and in an ovalbumin-dependent tolerization model in vivo, to limit IL-2 availability during Th1 differentiation, and to promote T cell–dependent colitis.^74–76^ Given that most autoimmune diseases, and AIH in particular, are characterized by expanded Th1 and IFN-γ producing CD8⁺ T cells,^21,45,57,66^ our data are consistent with a model in which the balance of IL-27 activity is tipped toward a pro-inflammatory role sustaining cytotoxic effector differentiation rather than tolerance in the chronically inflamed liver.

Unlike the PD-1⁺ TOX⁺ CD8⁺ T cells of cancer and chronic viral infection which are progressively rendered dysfunctional, cells in AIH livers produced multiple effector molecules, including granzymes, IFNγ, TNFα, and chemokines. This divergence may reflect not only the abundance of the progenitor exhausted T cell pools in the lymphoid tissues and liver but also the availability of niche signals such as IL-27 and IL-15 that are limiting in the classical exhaustion setting, and/or differences in TCR signal strength arising from a weaker affinity of self-antigens than viral antigens, either of which could tip a shared PD-1⁺TOX⁺ program away from dysfunction and toward sustained effector output. While, the relative contribution of various granzymes (GZMA/B/K) produced by the CD8^+^ T cells to hepatocyte damage in AIH still remains to be defined, we show that at least one predominant effector molecule, IFNγ, promoted inflammatory macrophage accumulation and thereby keeps the inflammatory circuit engaged: in vivo, loss of the IFNγR abrogated nearly all signs of liver pathology, indicating a strong requirement for IFNγ in initiating or propagating hepatic inflammation. We therefore favor a model in which accumulation of undigested DNA/RNA triggers the recruitment and activation of IFNγ producing innate lymphocytes in the liver, which then prime myeloid cells to more vigorously respond to TLR ligands,^77^ In this case, TLR9 detection of DNA. TLR9 signals in turn lead to accumulation of MoMacs that produce IL27 to promote T cell activation and survival. Together, these data identify macrophage activation pathways (e.g., TLR9 or JAK signaling) and IL-27/IL-27R axis as attractive therapeutic targets. Together, these findings resolve how autoreactive T cells remain functional under chronic antigen exposure and map a macrophage-centered circuit for selective intervention in AIH and beyond.

## ACKNOWLEDGEMENTS

We greatly appreciate input from our UMass colleagues: Drs. Michelle Kelliher, Pranoti Mandrekar, Milena Bogunovic, Lawrence Stern, and Kate Fitzgerald for overall experimental design and technical details, Dr. Zhaozhao Jiang for technical support on the preparation of RNAseq samples and initial bulk RNAseq analysis, and Drs. Alan Mullen and Cheng Sun for advice on the qPCR analysis of liver samples. We especially thank Dr. Maria Serena Longhi (BIDMC) for providing serum samples from AIH patients for ANA staining pattern evaluation, Dr. Zhijian Chen (UTSW) for providing liver tissues from Dnase2*^−/−^* Sting^Gt/Gt^ mice, and Dr. Jessica Hamerman (Benaroya Institute) for advice on hepatic immune cell isolation. We would like to thank Tim Nelson for helping with scRNAseq sample preparation. Melissa Strauss for performing ANA staining with human AIH serum samples. We would also thank Sharon Submaranian, Sebenele Lukhele, Sruthi Takillapati, and Stephanie Moses for genotyping and maintaining the mouse colony. These studies received invaluable assistance from the following UMass Chan Core Facilities: Animal Medicine, Flow Cytometry, and Morphology. Illustrations were created with BioRender.com. This project has been supported by NIH grant R37 AI155901 and American Liver Foundation.

## MATERIALS AND METHODS

### Key Resources Table

| REAGENT or RESOURCE | SOURCE | IDENTIFIER |
| --- | --- | --- |
| Antibodies |  |  |
| BV650 anti-mouse CD45 (30-F11) | Biolegend | Cat# 103151 |
| RB613 anti-mouse CD45(30-F11) | BD Pharmingen | Cat# 571489 |
| BUV805 anti-mouse CD11b (M1/70) | Invitrogen | Cat# 368-0112-82 |
| BV421 anti-mouse Ly6G (1A8) | Biolegend | Cat# 127628 |
| Pacific Blue anti-mouse I-A/I-E (M5/114.15.2) | Biolegend | Cat# 107620 |
| BV605 anti-mouse CD11c (N418) | Biolegend | Cat# 117331 |
| FITC anti-mouse Ly6C (HK1.4) | Biolegend | Cat# 128006 |
| BV570 anti-mouse Ly6C (HK1.4) | Biolegend | Cat# 128030 |
| PE-Dazzle594 anti-mouse CD43 (S11) | Biolegend | Cat# 143218 |
| PE anti-mouse TIM4 (F31-5G3) | Biolegend | Cat# 129906 |
| RB705 anti-mouse CD3 (17A2) | BD Pharmingen | Cat# 757248 |
| APC-Cy7 anti-mouse F4/80 (BM8) | Biolegend | Cat# 123114 |
| PE anti-mouse CD319 (4G2) | Biolegend | Cat# 152006 |
| PE-Cy5 anti-mouse CD64 (X54-5/7.1) | Biolegend | Cat# 139332 |
| APC anti-mouse CLEC2 (17D9) | Biolegend | Cat# 146106 |
| PE-Cy7 anti-mouse VISIG4 (NLA14) | Invitrogen | Cat# 25-5752-82 |
| APC-Fire810 anti-mouse CD19 (6D5) | Biolegend | Cat# 115578 |
| AF700 anti-mouse H2-Kb (AF6-88.5) | Biolegend | Cat# 116522 |
| BUV395 anti-mouse CD8a (53-6.7) | Invitrogen | Cat# 363-0081-82 |
| RB670 anti-mouse CD8a (53-6.7) | BD Pharmingen | Cat# 571948 |
| AF700 anti-mouse CD8b (YTS156.7.7) | Biolegend | Cat# 126618 |
| BUV661 anti-mouse CD4 (RM4-4) | BD Pharmingen | Cat# 612974 |
| BUV737 anti-mouse TCRb (H57-597) | Invitrogen | Cat# 367-5961-82 |
| PE-Cy7 anti-mouse TCRg/d (GL3) | Biolegend | Cat# 118124 |
| BV605 anti-mouse/human CD44 (IM7) | Biolegend | Cat# 103047 |
| FITC anti-mouse DX5 (DX5) | Biolegend | Cat# 108906 |
| PE anti-mouse NK1.1 (PK136) | BD Pharmingen | Cat# 557391 |
| PE-Dazzle594 anti-mouse CXCR6 (SA051D1) | Biolegend | Cat# 151117 |
| BV711 anti-mouse CXCR6 (SA051D1) | Biolegend | Cat# 151111 |
| PE-Cy7 anti-mouse CD49a (HMa1) | Biolegend | Cat# 142608 |
| BV750 anti-mouse CD49a (Ha31/8) | BD Pharmingen | Cat# 746854 |
| RB780 anti-mouse CD62L (MEL-14) | BD Pharmingen | Cat# 569209 |
| APC-eFluor780 anti-mouse CD62L (MEL-14) | Biolegend | Cat# 100476 |
| BUV496 anti-mouse PD-1(29F.1A12) | BD Pharmingen | Cat# 568598 |
| BUV563 anti-mouse KLRG1 (2F1) | Invitrogen | Cat# 365-5893-82 |
| BV480 anti-mouse SLAMF6 (13G3) | BD Pharmingen | Cat# 746534 |
| PE anti-mouse IL27Ra (W16125D) | Biolegend | Cat# 159004 |
| PE anti-mouse GzmA (3G8.5) | Biolegend | Cat# 149704 |
| PE-Cy5 anti-mouse GzmB (QA16A02) | Biolegend | Cat# 372226 |
| BV650 anti-mouse TNFa (MP6-XT22) | Biolegend | Cat# 506333 |
| BUV805 anti-mouse IFNgamma (XMG1.2) | Invitrogen | Cat# 368-7311-82 |
| PE-Cy7 anti-mouse TCF-7 | Cell Signaling | Cat# C6309 |
| efluor660 anti-mouse TOX (TXRX10) | Invitrogen | Cat# 50-6502-82 |
| FITC anti-mouse Ki-67 (SolA15) | Invitrogen | Cat# 11-5698-82 |
| BV421 anti-T-bet (4B10) | Biolegend | Cat# 644832 |
| RB613 anti-mouse RORgt (Q31-378) | BD Pharmingen | Cat# 571115 |
| AF700 anti-mouse/rat Foxp3 (FJK-16s) | Invitrogen | Cat# 56-5773-82 |
| AF700 anti-mouse CD45.1 (A20) | Biolegend | Cat# 110724 |
| APC-Cy7 anti-mouse CD45.1 (A20) | Biolegend | Cat# 110716 |
| BV750 anti-mouse CD45.2 (104) | Biolegend | Cat# 109857 |
| PE anti-mouse CD289 TLR9 (S18025A) | Biolegend | Cat# 159104 |
| APC anti-mouse CD289 (TLR9) (S18025A) | Biolegend | Cat# 159106 |
| Mouse LYVE-1 antibody | R&D Systems | Cat# AF2125 |
| Anti-mouse Cytokeratin 19 (EP1580Y) | Abcam | Cat# ab52625 |
| Donkey anti-goat AF647 | Invitrogen | Cat# A-21447 |
| Donkey anti-rabbit AF488 | Invitrogen | Cat# A-21206 |
| DyLight 488-coupled goat anti-mouse IgG Ab | Biolegend | Cat# Poly4053 |
| AffiniPure Goat Anti-Mouse IgG Fcgamma | Jackson ImmunoResearch | Cat# 115-005-205 |
| Anti-Mouse IgG (Fc-specific)- Peroxidase | Sigma Aldrich | Cat# A0168 – 1ML |
| Goat anti-mouse IgG2c | Southern Biotech | Cat# 1077-01 |
| Biotin Mouse Anti-Mouse IgG2a[b] (5.7) | BD Pharmingen | Cat# 555304 |
| Purified NA/LE Hamster Anti-Mouse CD3e | BD Pharmingen | Cat# 553057 |
| Purified NA/LE Hamster Anti-Mouse CD28 | BD Pharmingen | Cat# 553294 |
| Chemicals, peptides, and recombinant proteins |  |  |
| 10% neutral buffered formalin | Fisher HealthCare | Cat# 23-245685 |
| Z-Fix | Anatech | Cat# 175 |
| Xylene | Fisher | Cat# 9990501 |
| Tris-EDTA pH 9.0 antigen retrieval buffer (10x) | Biolegend | Cat# 422703 |
| Triton X-100 | Sigma Aldrich | Cat# X100 |
| Normal donkey serum | Abcam | Cat# ab7475 |
| DAPI | Thermo fisher | Cat# D1306 |
| Prolong diamond antifade mountant | Thermo fisher | Cat# P36970 |
| RPMI 1640 Medium | Thermo Fisher | Cat# 11875093 |
| StemSpan™ SFEM | Stemcell Technologies | Cat# 09600 |
| Percoll | Cytiva | Cat# 17-0891-01 |
| Ghost Dye Violet510 | Tonbo Bioscience | Cat# 130870T100 |
| Ionomycin | Cell Signaling | Cat# 9995S |
| PMA | MilliporeSigma | Cat# P1585 |
| BD GolgiPlug™ Protein Transport Inhibitor (Containing Brefeldin A) | BD Pharmingen | Cat# 555029 |
| Human IL-27 Recombinant Protein, PeproTech | Thermo Fisher | Cat# 200-38-10UG |
| Mouse IL-15 Recombinant Protein, PeproTech | Thermo Fisher | Cat# 210-15-10UG |
| ACK Lysing buffer | Thermo Fisher | Cat# A1049201 |
| FluoroFix buffer | Biolegend | Cat# 422101 |
| DNA/RNA Shield | Zymo Research | Cat# R1100-50 |
| Xenium Decoding Reagent Module A (Universal) | 10X Genomics | Cat# 1000859 |
| Xenium Prime Decoding Reagent Module B - 5K | 10X Genomics | Cat# 1000722 |
| Xenium Cell Segmentation Staining Reagents | 10X Genomics | Cat# 1000661 |
| Critical commercial assays |  |  |
| Alanine Transaminase Activity Assay Kit | Abcam | Cat# ab105134 |
| Liver Dissociation Kit, mouse | Miltenyi | Cat# 130-105-807 |
| Lineage Cell Depletion Kit, mouse | Miltenyi | Cat# 130-110-470 |
| Zombie NIR Fixable Viability Kit | Biolegend | Cat# 423105 |
| Fixation/permeabilization solution kit | BD Pharmingen | Cat# 554714 |
| FoxP3/transcription factor staining buffer kit | Invitrogen | Cat# 00-5523-00 |
| MojoSort mouse CD8 T cell isolation kit | Biolegend | Cat# 480008 |
| TMB substrate reagent set | BD Pharmingen | Cat# 555214 |
| RNeasy mini kit | Qiagen | Cat# 79216 |
| iScript gDNA clear cDNA synthesis kit | Bio-rad | Cat# 1725035 |
| Taqman fast advanced master mix | Life Technologies | Cat# 4444556 |
| RNeasy plus micro kit | Qiagen | Cat# 74034 |
| Illumina stranded mRNA prep, Ligation | Illumina | Cat# 20040532 |
| IDT for Illumina RNA UD Indexes Set A, Ligation | Illumina | Cat# 20040553 |
| SPRIselect beads | Beckman Coulter | Cat# B23317 |
| Qubit dsDNA quantification assay kit | Invitrogen | Cat# Q32851 |
| Nextseq 2000 P2 reagent kits | Illumina | Cat# 20046813 |
| Single Cell 5' Library & Gel Bead Kit | 10X Genomics | Cat# 1000165 |
| Chromium Single Cell V(D)J Enrichment Kit, mouse T cells | 10x Genomics | Cat# 1000071 |
| Xenium Prime Sample Prep Reagents (2 reactions) kit | 10X Genomics | Cat# 1000720 |
| Xenium Prime 5K Mouse Pan Tissue & Pathways Panel (2 rxns) | 10X Genomics | Cat# 1000725 |
| Xenium Decoding Consumables v2 |  | Cat# 1000726 |
| Nucleofector solution set P3 | LONZA | Cat# PBP3-00675 |
| Gene Knockout Kit (mouse) - Il27 | EditCo | https://store.editco.bio/products/gene-knockout-kits |
| Gene Knockout Kit (mouse) - Il27ra | EditCo | https://store.editco.bio/products/gene-knockout-kits |

| Experimental models: Organisms/strains |  |  |
| --- | --- | --- |
| Mouse: C57BL/6-Tlr9em1.1Ldm/J (TLR9 KO) | Jackson Laboratories | Strain# 034449 |
| Mouse: B6.129S4-Ifngtm3.1Lky/J (GREAT) | Jackson Laboratories | Strain# 017581 |
| Mouse: B6.SJL-Ptprca Pepcb/BoyJ (CD45.1) | Jackson Laboratories | Strain# 002014 |
| Mouse: B6.129S7-Rag1tm1Mom/J (RAG1 KO) | Jackson Laboratories | Strain# 002216 |
| Mouse: Dnase2-/- Ifnar-/- | Our Lab | N/A |
| Mouse: Dnase2-/- Ifnar-/- Tlr7-/- | Our Lab | N/A |
| Mouse: Dnase2-/- Ifnar-/- Ifngr-/- | Our Lab | N/A |
| Mouse: Dnase2-/- StingGt/Gt | Chen Lab | N/A |
| Mouse: Un93b1-/- | Latz Lab | N/A |
| Oligonucleotides |  |  |
| Gapdh (Mm99999915_g1) | Life Technologies | Cat# 4331182 |
| Col1a1(Mm00801666_g1) | Life Technologies | Cat# 4331182 |
| Tnfa (Mm00443258_m1) | Life Technologies | Cat# 4331182 |
| Ifng (Mm01168134_m1) | Life Technologies | Cat# 4331182 |
| Cxcl10 (Mm00445235_m1) | Life Technologies | Cat# 4331182 |
| Software and algorithms |  |  |
| FlowJo 10.10.0 | Flowjo | https://www.flowjo.com |
| Graphpad prism 10.2.1 | GraphPad Software | https://www.graphpad.com/ |
| CFX manager 3.1 | Bio-rad | N/A |
| Qupath 0.7.0 | Bankhead et al.<sup>78</sup> | https://qupath.github.io/ |
| TissueFAXS Viewer 8.0.0 | Tissue Gnostics | https://tissuegnostics.com/products/tissuefaxs-platform |
| R 4.4.3 | CRAN | https://cran.r-project.org/mirrors.html |
| Rstudio | RStudio | https://rstudio.com/ |
| Seurat 5.4.0 | Hao, et al.<sup>79</sup> | https://satijalab.org/seurat/ |
| scRepertoire 2.2.1 | Yang et al.<sup>80</sup> | https://www.borch.dev/uploads/screpertoire/ |
| fGSEA 1.32.4 | Korotkevich et al.<sup>81</sup> | https://github.com/alserglab/fgsea |
| DESeq2 1.46.0 | Love et al.<sup>82</sup> | https://github.com/thelovelab/DESeq2 |
| Monocle3 1.4.27 | Trapnell et al.<sup>83</sup><br>Qiu et al.<sup>84</sup><br>Cao et al.<sup>85</sup> | cole-trapnell-lab.github.io/monocle3 |
| Proseg 3.2.0 | Jones et al.<sup>86</sup> | https://github.com/dcjones/proseg |
| Scanpy 1.11.5 | Wolf et al.<sup>87</sup> | https://scanpy.scverse.org/en/stable/ |
| nichenetr 2.2.1.1 | Browaeys et al.<sup>88</sup> | https://github.com/saeyslab/nichenetr |
| Others |  |  |
| gentleMACS Octo Dissociator with Heaters | Miltenyi Biotec | N/A |
| 5-Laser Cytex Aurora spectral analyzer | Cytex | N/A |
| Multi-Application MA900 cell sorter | Sony Biotechnology | N/A |
| CFX Opus 384 Real-Time PCR System | Bio-rad | Cat# 12011452 |
| 4D-Nucleofector Core Unit | Lonza | Cat# AAF-1003B |
| 4D-Nucleofector X Unit | Lonza | Cat# AAF-1003X |
| Nextseq 2000 sequencing system | Illumina | N/A |

### Mice

DNase II-gene targeted mice were kindly provided by Dr. S. Nagata and obtained through the Riken Institute. Dnase2^+/−^ x Ifnar^−/−^ (Het), Dnase2^−/−^ x Ifnar^−/−^ (DKO), Dnase2^−/−^ x Ifnar^−/−^ x Tlr7^−/−^ (TLR7 TKO), Dnase2^−/−^ x Ifnar^−/−^ x Ifngr^−/−^ (IFNgR TKO) mice have been described previously (14). Dnase2*^−/−^* Sting^Gt/Gt^ mice were kindly provided by Dr. Z. Chen (UT Southwestern Medical Center, Dallas, TX). Tlr9^-/-^ (Jax #034449), IFNγ reporter B6.129S4-Ifng^tm3.1Lky^/J (Jax# 017581), and B6 CD45.1 (Jax #002014) mice were obtained from Jackson Laboratory. Unc93b1^- /-^ mice were kindly provided by Dr. E. Latz. DKO mice were intercrossed with Tlr9^-/-^, Unc93b1^-/-^, B6.129S4-Ifng^tm3.1Lky^/J, and B6 CD45.1 mice to generate the corresponding triple KO lines, DKO X GREAT line, and CD45 allotype distinct line. Mice were euthanized by isoflurane. Serum was collected by cardiac puncture on euthanized animals. All animal procedures were approved and performed in accordance with the Institutional Animal Care and Use Committee at the University of Massachusetts Medical School.

### Liver Histology and quantification

The left lateral lobe of livers was removed, fixed with 10% phosphate buffered formalin (PBF) and embedded in paraffin. Hematoxylin and eosin (H&E), Masson’s trichrome, and TUNEL staining on paraffin-embedded liver sections were carried out by the Morphology Core (University of Massachusetts Chan Medical School, Worcester, MA). Sirius red staining was carried out by Applied Pathology Systems (Shrewsbury, MA). Slides were scanned at 20x using TissueFAXS platform (TissueGnostics) by Sanderson Center for Optical Experimentation (University of Massachusetts Chan Medical School, Worcester, MA). Percent of Sirius red positive area relative to scanned tissue area was quantified using Qupath.

### Immunofluorescence

FFPE embedded liver blocks were sectioned at 8uM for immunofluorescent staining targeting LYVE-1 and CK19. After deparaffinization and rehydration, liver sections were subjected to heat-induced antigen retrieval with Tris-EDTA pH 9.0 antigen retrieval buffer (Biolegend) in pressure cooker for 15 mins, permeabilized with 0.3% Triton-X (Sigma Aldrich) for 10 mins, and blocked with blocking buffer (10% normal goat serum and 1% bovine serum albumin in PBS) for 1h at RT. Sections were then incubated with primary antibodies overnight in 4C, followed with fluorophore-conjugated secondary antibodies for 1h at RT. After washing, sections were stained with DAPI (Thermo fisher) for 5 minutes at RT and mounted with Prolong diamond anti-fade mounting reagents (Thermo fisher). Stained liver slides were imaged with a TUNDER Imager (Leica). F4/80, CD4, and CD8 multiplex immunofluorescence staining was performed on FFPE embedded liver blocks at UCSD Biorepository and Tissue Technology Center as fee for service.

### qPCR analysis of liver tissues

A small piece of liver left lobe was homogenized in RLT buffer (QIAGEN). RNA was extracted using RNeasy Mini kit (QIAGEN). Complimentary DNA was reversely transcribed from total RNA using iScript gDNA clear cDNA synthesis kit (Bio-rad). Real-time PCR was performed with Taqman fast master mix (Life Technologies) and the following pre-designed probes: *Tnfa* (Mm00443258_m1), *Ifng* (Mm01168134_m1), *Cxcl10* (Mm00445235_m1), and *Gapdh* (Mm99999915_g1) (Life Technologies) on a Bio-rad CFXOpus 384 Real-time PCR system. Expression level of each target gene was normalized to *Gapdh* expression.

### Plasma alanine aminotransferase activity

Serum samples were collected from 3–4-month-old mice. Serum levels of alanine aminotransferase (ALT) activity were determined using the Alanine Transaminase Activity Assay Kit (Abcam) following manufacturer’s instruction.

### Serum Immunoglobulin and Cytokine Analysis

Serum IgG2a and IgG1 were measured by ELISA as previously described.^89^ Antibodies are listed in key resources table.

### Antinuclear antibody assay

Slides coated with Hep-2 cells (Kallestad) were stained with serum samples (1:50 dilution) collected from 3–4-month-old mice. Bound antibodies were detected with DyLight 488-coupled goat anti-mouse IgG Ab (Biolegend). Samples were imaged with a TUNDER Imager (Leica) at 40X.

### Immune cell isolation

To isolate immune cells from the liver for flow cytometry, livers were perfused with 10 ml ice-cold PBS through the portal vein, then homogenized using Liver Digestion Kit (Miltenyi) and GentleMACS Octo Dissociator (Miltenyi). Cell suspensions were passed through a 100uM cell strainer, washed with plain DMEM and PBS containing 2mM EDTA and 0.5% BSA, and centrifuged at 300g for 10 min. RBCs were removed with ACK lysing buffer (Gibco # A1049201).

To isolate liver immune cells for RNA sequencing, cell suspension derived from GentleMACS dissociation was further purified by Percoll gradient centrifugation. The cells were washed twice with PBS containing 2mM EDTA, 0.5% BSA, then resuspended in 33% Percoll (Cytiva). Cell suspensions were centrifuged at 800g for 20 min at room temperature with no brake. The supernatant containing liver parenchymal cell debris was discarded, RBCs in the cell pellet were lysed, and the remaining cells were resuspended in RPMI medium containing 10% fetal calf serum (FCS) for later use.

Spleens were mashed and filtered through 70uM cell strainers in Hanks’ Balanced Salt Solution (HBSS) + 2% FCS, followed by RBC lysing.

### Flow cytometry and cell sorting

#### Surface staining

Cell suspensions were first stained with Zombie NIR Fixable Viability kit (Biolegend) or Ghost Dye Violet 510 (Cytek) in PBS. Cells were then incubated with anti-CD16/31 (2.4G2) antibody to block Fc receptors and stained with fluorophore conjugated primary antibodies in PBS + 3% FCS at 4°C for 30 min. Antibodies were purchased from Biolegend, Tonbo Biosciences, eBiosciences, and Waters Biosciences and listed in key resources table. In some experiments, samples stained with only surface markers were fixed with FluoroFix buffer (Biolegend) prior to analyzed by flow cytometer.

#### Intracellular staining

Following surface marker staining, cells were fixed and permed using the FoxP3/Transcription factor staining buffer kit (Invitrogen #00-5523-00) according to manufacturer’s protocol. To detect cytokine production from T cells, percoll purified immune cells were restimulated with PMA and Ionomycin in RPMI + 10% FCS containing Golgiplug (BD pharmingen) for 4h at 37°C. For TLR9 detection, extracellularly stained cells were permed with BD Cytofix/Cytoperm fixation/permeabilization solution (BD pharmingen) for 20 min at 4°C, washed, and stained with fluorophore conjugated anti-TLR9 antibodies (Biolegend) in BD Perm/Wash Buffer (BD pharmingen) for 45 min at 4°C.

#### Data acquisition

The absolute cell counts were determined using Precision Count Beads (Biolegend #424902) and expressed as the number of cells per gram of tissue. Data were acquired on a 5-laser Aurora Cytek and analyzed with FlowJo software v10.10.0 (Tree Star). For bulk and single cell RNAseq, cells were sorted with FACSAria (Waters Biosciences) or SONY MA900 cell sorter (Sony Biotechnology).

### Generation of radiation chimeras, CRISPR knockout

Lethally irradiated (8.5Gy) 7-9-week-old Het and DKO mice mice were reconstituted with 10^7^ bone marrow cells from sex and age matched mice by i.v. injection. Bone marrow cells were harvested by flushing tibia and femurs of the donor mice. Recipient mice were then maintained on sulfatrim water. Chimerism and disease severity were assessed 9 weeks post reconstitution. The extent of reconstitution was determined by flow cytometry based on CD45 allelic markers.

For IL27 and IL27R knockout, bone marrow cells from DKO mice were first subjected to lineage cell depletion using mouse lineage cell depletion kit (Miltenyi) to enrich for stem cells following manufacturer protocol. CRISPR-Cas9-mediated knockout of *Il27* or *Il27ra* in bone marrow stem cells was performed by electroporation using lonza 4D-nucleofector system (Lonza Biosciences) with mouse gene-specific gene knockout kits (EditCo). Cells electroporated with non-targeting guide RNAs designed by EditCo served as negative controls. Cells post electroporation were immediately transferred to Stemspan media (Cell Technology) and rested for 30 mins at 37C before resuspended in PBS for transplant. Lethally irradiated RAG TKO hosts (8 Gy) were intravenously injected with 3-5 x 10^5^ stem cells. Chimerism and disease severity were assessed 9 weeks post reconstitution.

### In vitro CD8^+^ T cell culture

CD8^+^ T cells were isolated from WT spleen using MojoSort mouse CD8 T cell isolation kit (Biolegend). Cells were initially stimulated with 5ug/mL plate-bound anti-CD3 and anti-CD28 (BD Biosciences) for 72h. Cells were then harvested and treated with 50ng/ml IL-2, IL-15, or IL-27 (Thermo fisher) in complete RPMI for additional 24h before collected for bulk RNAseq.

### Bulk RNA sequencing and analysis

Total RNA from sorted monocyte/macrophages were isolated using RNeasy plus micro kit (QIAGEN) following manufacturer’s protocol. cDNA library was prepared using the Illumina Stranded mRNA Prep kit including indexes (Illumina). Libraries were purified using SPRIselect beads (Beckman Coulter). Library quality and size distribution were assessed using Bioanalyzer by UMass Molecular Biology core, and the concentration was determined using Qubit dsDNA quantification assay kit (Thermo fisher). Pooled libraries were sequenced in-house on an Illumina NextSeq 2000 using a P2 reagent kit. Sequencing data was processed as described previously using an RNAseq pipeline (DolphinNext/ViaFoundry) to align and quantify mRNA transcripts as expected counts (RSEM). Resulting RSEM values were processed by DESeq2 package for identification of differentially expressed genes (padj < 0.01 and fold change >2). Principal component analysis was performed on the top 500 most variable genes.

For bulk RNAseq on in vitro cultured T cells, cells were lysed in DNA/RNA Shield (Zymo Research) and sent to Plasmidsaurus for cDNA library preparation and 3’ end-counting RNA-Seq using the Illumina platform. Raw count data were normalized, and differential gene expression analysis was performed using the DESeq2 package.

### Single-cell RNA/TCR sequencing

Single cell suspensions were prepared from liver and spleens as described above. CD3^-^ fraction from livers from 2 Het and 2 DKO mice were sorted and pooled within genotype for scRNAseq; CD3^+^ fraction from livers and spleens of Het and DKO mice were sorted for paired scRNA/TCRseq. Libraries were generated as previous described.^90^ Approximately 20,000 cells per sample were loaded onto a 10x Chromium Chip G and 5′ gene expression libraries were prepared with the Chromium Single-Cell 5′ Library and Gel Bead Kit (10x Genomics) according to manufacturer’s instructions. T cell receptor libraries were prepared from the CD3+ samples with the mouse T cell Single Cell V(D)J Enrichment Kit (10x Genomics). Library quality was assessed using a TapeStation (Agilent) and sequenced by NovaSeq6000 (Illumina) to obtain a minimum of 20,000 paired-end reads per cell. Single-cell TCR libraries were multiplexed and sequenced by NovaSeq6000 (Illumina) to obtain a minimum of 5000 paired-end reads per cell. The sequencing specifications were set according to the manufacturer’s specification (10x Genomics).

### Processing of scRNA/TCR seq data

Reads were aligned and quantified with Cell Ranger v9.0.1 against refdata-cellranger-GRCm39-2024-A (gene expression) and refdata-cellranger-vdj-GRCm38-alts-ensembl-7.1.0 (V(D)J). Cell Ranger output was analyzed in Seurat v5.5.1 (R v4.4.3): after normalization and scaling, cells with 200–5,000 detected genes and under 10% mitochondrial content were retained. Libraries were integrated with Harmony and principal components used for UMAP and clustering. Cluster identities were assigned from differentially expressed markers (FindAllMarkers, Seurat).

For processing of scTCRseq data, TCR contigs from the Cell Ranger filtered contig annotations output were integrated with the Seurat object by matching cell barcodes using scRepertoire. CD8⁺ T cells with at least one productive TCRα- or TCRβ-chain CDR3 nucleotide sequence were retained for repertoire analysis.

The developmental trajectory of activated CD8 T cells was inferred with Monocle v3.0. Gene set enrichment for the monocyte/macrophage cluster was performed with fgsea v1.32.2 against Gene Ontology biological process terms (GO.db v3.20.0), with genes preranked by log2 fold change. Antigen-processing and presentation scores were calculated using UCell from genes annotated to GO:0019882 and GO:0002486 (UCell v.2.10.1). UCell scores were derived from the within-cell expression ranks of signature genes using the Mann–Whitney U statistic, with higher scores indicating stronger enrichment of the corresponding gene signature.

### Xenium in situ spatial transcriptomics

Livers from 2-month-old DKO and Het mice were fixed in 10% neutral buffered formalin for 24h at 4 °C, paraffin-embedded, and sectioned at 10uM. Liver sections were processed with the Xenium Prime Sample Prep Reagents kit (10x Genomics), and Xenium Cell Segmentation Staining Reagents (10x Genomics) according to manufacturer’s instructions and run on a Xenium Analyzer (10x Genomics). Liver sections from both genotypes were mounted on the same Xenium slide and processed in a single run. 5056 genes were targeted including 5,006 genes from Xenium Mouse 5K Pan Tissue and Pathways base panel plus 50 custom targets (Table S1).

### Segmentation, quality control and annotation

Cells were segmented de novo from transcript coordinates with Proseg (v3.2.0), which models cell shape and local expression and flags ambient background to reduce hepatocyte-marker bleed into non-hepatocyte cells. 66,047 cells from DKO and 76,381 cell Het were recovered. Cells with ≥10 transcripts and ≥5 genes were retained (n = 142,276), median-normalized, log-transformed and clustered per section using the top 50 Principal components, and Leiden resolution of 2.0 with Scanpy package (v1.11.5). Clusters were designated by z-scored marker gene expression, then immune and mesenchymal cells refined per cell on raw counts. Sections were concatenated and re-embedded without batch correction.

### Processing of Xenium data

Macrophages were scored on a Kupffer axis based on mean Cd5l, Vsig4 expression and displayed continuously between the 2nd and 98th percentiles. Hepatocytes were placed on a porto-central axis (mean Glul, Cyp2e1, Cyp1a2, Cyp2a5, Axin2, Lgr5 expression minus mean Hal, Ass1, Arg1, Gls2 expression) and cut into zones at Het quartiles; the cell-type atlas instead shades hepatocytes in three equal groups for orientation. The distance between CD8⁺ T cell and the nearest macrophage, dendritic cells, CD4⁺ T cell, B cells, and NK cells within the same tissue section was computed using a k-d tree nearest-neighbor search. Distance distributions were visualized using Gaussian kernel density estimates with a bandwidth of 8 µm. Chemokine association is transcript density (taken at raw coordinates without cell assignment) within 20 µm of a CD8 T cell over tissue-wide density. For visualization, 375 x 375 µm regions were selected from the chemokine expression map. Transcript counts for Cxcl9, Cxcl10, and Cxcl16 were aggregated in 25 x 25 µm bins respectively and normalized to total transcript count within the tissue. Normalized expression from the three chemokines were summed and smoothed to generate a combined chemokine-density map. Three chemokine-rich windows focused on local maxima of this map, whereas contrast windows were selected from regions with chemokine signals below the 35^th^ percentile.

### NicheNet analysis

Putative ligand-receptor interactions between CD8^+^ T cells with other populations from spatial transcriptomic and scRNAseq dataset was inferred using nichenetr package (v2.2.1.1) and the NicheNet v2 mouse prior model (Zenodo 7074291). Ligand activity was quantified using the area under the precision-recall curve (AUPR) to assess how each ligand’s predicted targets recovered the 1489 genes upregulated in DKO CD8+ T cells (log₂ fold change > 0.25 and adjusted *P* < 0.05) from a background of 5045 genes. A total of 151 candidate ligands were evaluated and prioritized based on ligand differential expression, receptor differential expression, scaled ligand activity, and receptor expression. For each ligand, target genes with predicted regulatory-potential scores above 5% of that ligand’s highest-scoring target were retained.

### Processing of published RNA-seq data

Public scRNAseq dataset of PBMC from AIH and healthy donors was obtained from GEO: GSE216064.^58^ T and NK cells were subclustered and annotated based on differentially expressed markers (FindAllMarkers, Seurat). Single-nucleus RNA-seq data from human liver biopsies, including the processed data and previously defined CD8⁺ T-cell and myeloid cluster annotations, were obtained from Sherman et al.^57^ Gene signatures for AIH-enriched myeloid clusters were defined using marker genes meeting the indicated thresholds (AUROC ≥ 0.65, fold change in percentage of expressing cells (%FC) ≥ 1.5, and log2FC > 0.8 for cluster 5, or log2FC > 1 for cluster 0 and 7). Selected human genes were mapped to their mouse orthologs, and module scores were calculated in the mouse scRNA-seq dataset using the Seurat AddModuleScore function. Scores were visualized on the UMAP of mouse monocyte/macrophage subsets.

### Statistical Analysis

All statistical analyses were performed in Prism V.10 (GraphPad Software). Sample size in each experiment is detailed in the figure legend, where n = number of mice. Normality of the data distribution was examined by Shapiro-Wilk tests. The statistical significance of differences between two groups was determined by Welch T test for normally distributed data, and by nonparametric tests for data that are non-normal distributed. One-way ANOVA was used when comparing more than two groups. Differences were considered significant if P < 0.05, and the following represents the level of significance: ****P < 0.0001, ***P < 0.001, **P < 0.01, and *P < 0.05.

## Author Contributions

Conceptualization: K.H., A.M.R, and V.S.K

Data Curation: K.H., A.S.V.B., S.G.B., D.M.S., M.S.S., V.S.K.

Methodology: K.H., K.N., K.M.G., A.M.R., V.S.K

Validation: K.H., A.S.V.B., S.G.B.

Formal Analysis: K.H., S.G.B., A.S.V.B., D.M.S., M.S.S., V.S.K.

Investigation: K.H., A.S.V.B., S.G.B., P.L.K., S.V.K., K.C., D.M.S., M.S.S., V.S.K.

Writing – original draft: K.H., A.M.R., V.S.K

Writing – review & editing: K.H, S.G.B., A.M.R., V.S.K Visualization: K.H., S.G.B.

Supervision: A.M.R., V.S.K

Project Administration: K.H., A.M.R., V.S.K Resources: A.M.R., V.S.K

Funding Acquisition: V.S.K, A.M.R.

---

**Supplemental Figure 1.**
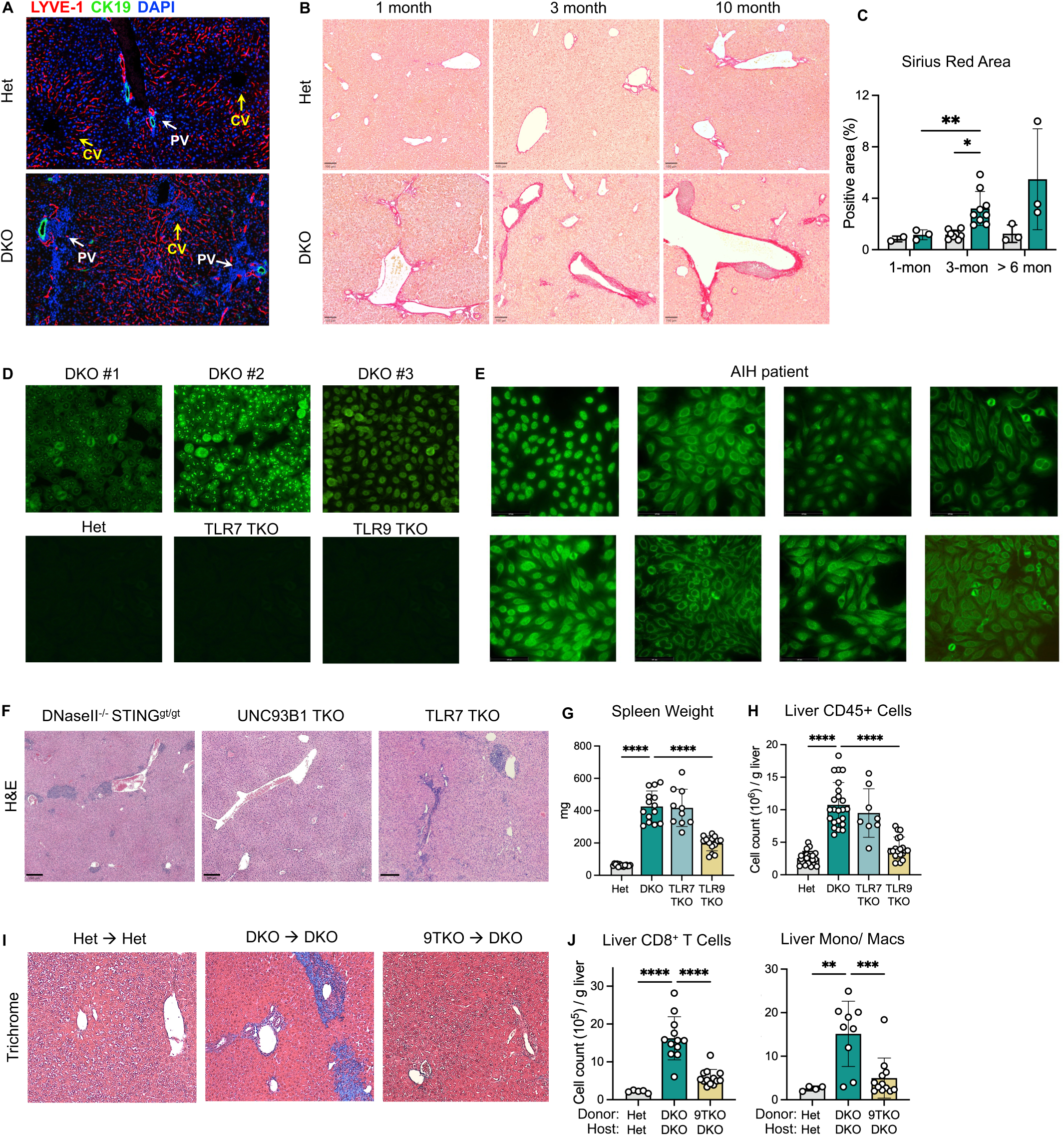
(A) Representative immunofluorescence images of liver section from Het, DKO, TLR7 TKO, and TLR9 TKO mice showing bile duct (CK19^+^ green), sinusoidal endothelial cells (LVYE-1^+^, red), and cell nucleus (DAPI, blue). (B) Representative Sirius red staining (scale bar, 100um) of liver sections from Het and DKO mice at indicated age. (C) Quantification of Sirius red positive liver area. (D) Representative images of HEp-2 cells stained with sera from 3-4-month-old mice to reveal ANAs. (E) Representative images of HEp-2 cells stained with sera from AIH patients to reveal different ANA staining patterns (n = 8). (F) Representative H&E staining of liver sections from UNC93B1 TKO, DNaseII^-/-^ STING^gt/gt^ mice, and TLR7 TKO mice. (G) Spleen weight of 3-4 month-old mice of indicated genotype (n = 10-14 per group). (H) Absolute numbers of CD45^+^ immune cells in the livers of 3-4 month-old mice (n = 7-25 per group). (I) Representative Masson’s trichrome staining (magnification,10X) of liver sections from chimeras 8-week post bone marrow transplant. (J) Absolute numbers of indicated immune population in the livers of chimeras 8-week post bone marrow transplant (n = 9-14 per group).. Data are pooled from at least 3 independent experiments and represented as mean ± SEM. *p < 0.05, **p < 0.01, ***p < 0.001, and ****p < 0.0001

**Supplemental Figure 2.**
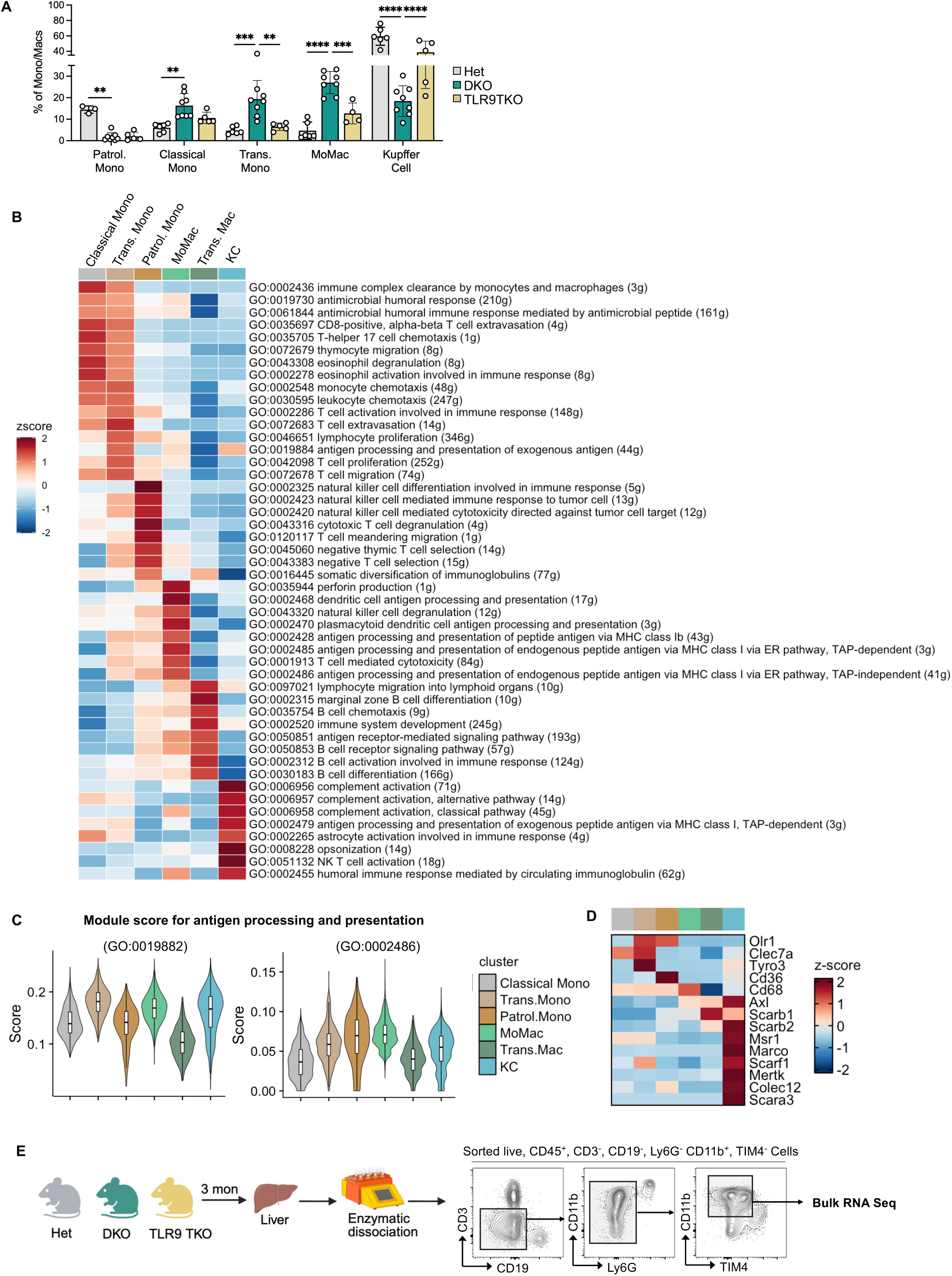
(A) Frequencies of indicated monocyte/macrophage subsets identified in figure 2D in Het, DKO, and TLR9 TKO livers (n = 5-8 per group). Data is pooled from 2 independent experiments and represented as mean ± SEM. *p < 0.05, **p < 0.01, ***p < 0.001, and ****p < 0.0001 (B) Heatmap showing enriched gene ontology (GO) terms in each monocyte/macrophage subset. (C) Antigen-presentation signature scores across monocyte and macrophage subsets based on genes in GO pathways: antigen processing and presentation of endogenous peptide antigen via MHC class I via ER pathway, TAP-independent (GO:0002486), and antigen processing and presentation (GO:0019882). (D) Heatmap showing RNA expression of scavenger receptor genes in indicated subsets across monocyte/macrophage subsets. (E) Gating strategy used to sort monocyte-lineage cells for bulk RNA sequencing in figure 3I.

**Supplemental Figure 3.**
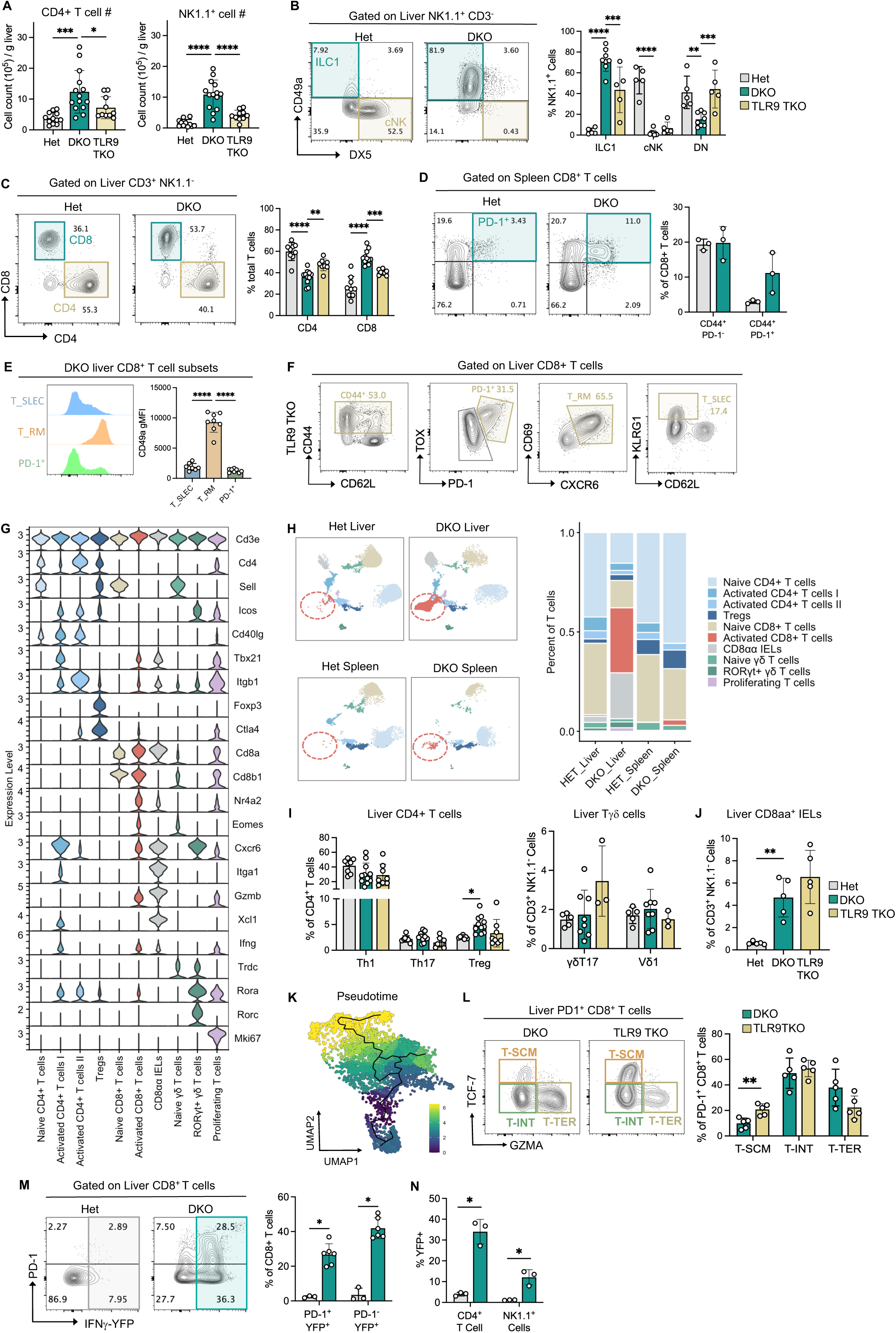
(A) Absolute number of CD4^+^ T cells, and NK1.1^+^ cells from livers of Het, DKO, and TLR9 TKO mice. (B) Representative flow cytometry plots and frequencies of ILC1, conventional NK (cNK), and DX5 CD49a double negative (DN) cells among liver CD3^-^ NK1.1^+^ cells ( n=5-8 per group). (C) Representative flow cytometry plots and frequencies of CD4+ and CD8+ T cells among liver CD3^+^ NK1.1^−^ cells (n = 9-14 per group). (D) Representative flow cytometry plots and frequencies of CD44^+^ PD-1^+^, and CD44^+^ PD-1^-^ T cells among spleen CD8^+^ T cells (n = 3 per group). (E) Representative histograms and quantification of CD49a expression in the indicated DKO liver CD8+ T cell subsets (n = 8). Expression is reported as gMFI. (F) Representative flow cytometry plots showing CD8+ T cell subsets identified in figure 3B in livers of TLR9 TKO mice. (G) RNA expression of genes defining the T cell subsets in figure 3C. (H) UMAP plots and relative abundances of T cell subsets from the livers and spleens of Het and DKO mice. (I) Frequencies of Th1 (T-bet^+^), Th17 (RORγT^+^), and regulatory T cells (Treg) (FoxP3^+^) among liver CD4+ T cells ( n= 8-11 per group); frequencies of γδT17 cells (RORγT^+^, IL17^+^) and Vδ1 (T-bet^+^) T cells among liver γδ T cells ( n= 3-8 per group). (J) Frequencies of liver CD8αα+ intraepithelial lymphocytes (IELs) among CD3^+^ NK1.1^−^ cells ( n= 5 per group). (K) Pseudotime trajectory analysis of activated liver CD8^+^ T cells by monocle 3. (L) Representative flow cytometry plots and frequencies of T_SCM, T_INT, and T_TER cells among liver CD44^+^ PD-1^+^ CD8^+^ T cells from DKO and TLR9 TKO mice (n = 5 per group). (M) Representative flow cytometry plots and frequencies YFP^+^ PD-1^+^ and YFP^+^ PD-1^-^ cells among liver CD8^+^ T cells in IFNγ reporter (GREAT) Het and DKO mice (n = 3-6 per group). (N) Frequencies of YFP^+^ cells among liver CD4+ T cells and NK1.1+ cells in IFNγ reporter (GREAT) Het and DKO mice (n = 3 per group). Data are pooled from at least 2 independent experiments and are presented as mean ± SEM. *p < 0.05, **p < 0.01, ***p < 0.001, and ****p < 0.0001.

**Supplemental Figure 4.**
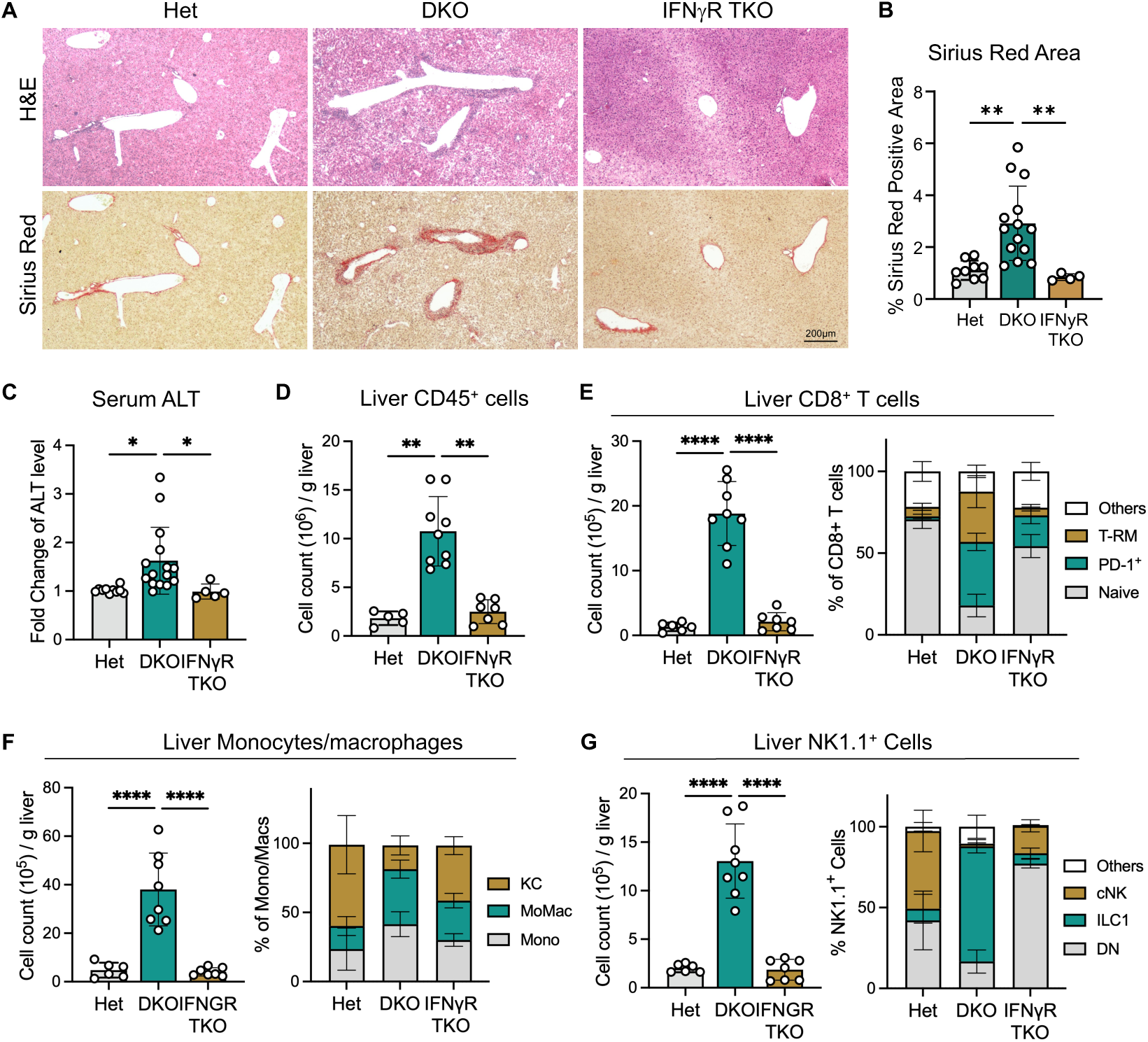
(A) Representative H&E, and Sirius Red staining (scale bars, 200um) of liver sections from 3-4-month-old mice. (B) Quantification of Sirius red positive liver area (n = 4-14 per group). (C) Serum ALT level relative to Het (n = 5-15 per group). (D) Absolute numbers of CD45^+^ immune cells in the livers (n = 5-9 per group). (E) Absolute numbers and subset distribution of CD8^+^ T cells in the livers (n = 6-8 per group). (F) Absolute numbers and subset distribution of monocyte/macrophage populations in the livers (n = 6-8 per group). (G) Absolute numbers and subset distribution of NK1.1^+^ cells in the livers (n = 6-8 per group). Data are pooled from at least 3 independent experiments and represented as mean ± SEM. *p < 0.05, **p < 0.01, ***p < 0.001, and ****p < 0.0001

**Supplemental Figure 5.**
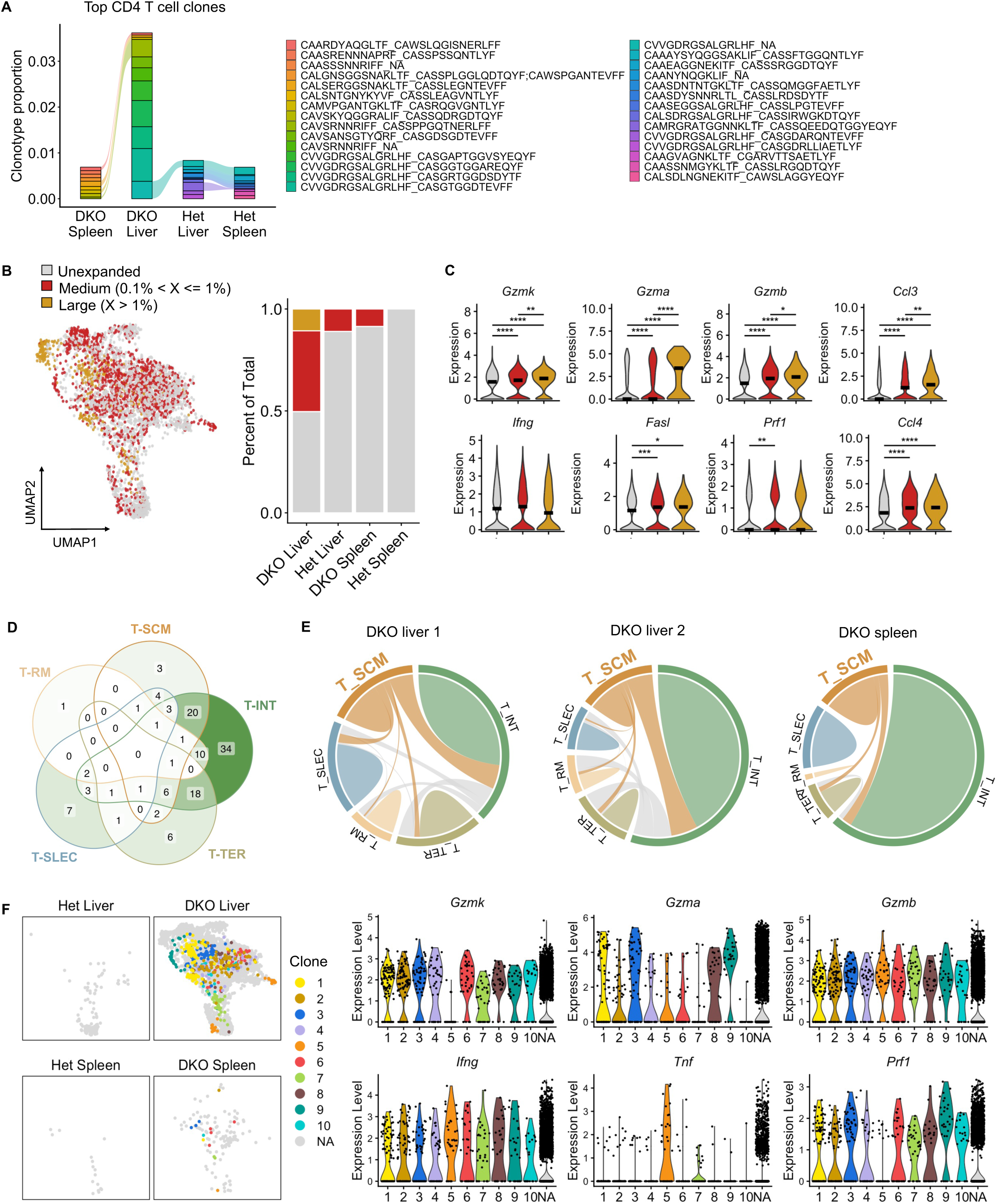
(A) Top expanded CD4^+^ T cell clonotypes across tissues, with each color representing one clone. (B) UMAP plot of activated CD8^+^ T cells showing TCR clone size. Bar graph summarizes relative abundance of large (yellow), medium (red), and unexpanded (grey) clones within activated CD8^+^ T cells in the livers and spleens of Het and DKO mice. (C) RNA expression of cytotoxic molecule across unexpanded, medium, and large TCR clones. *p < 0.05, **p < 0.01, ***p < 0.001, and ****p < 0.0001. (D) Venn diagram showing the overlap of expanded TCR clonotypes (clone size > 0.1%) among indicated CD8+ T cell subsets. Numbers indicates clonotype counts within each unique or shared region. (E) Chord diagrams showing shared TCR clonotypes among activated CD8+ T cell subsets in two individual DKO liver samples and pooled DKO spleen sample. (F) UMAP plots of activated CD8^+^ T cells showing the tissue distribution of the top 10 shared TCR clonotypes identified in Figure 4I; and RNA expression of indicated cytotoxic molecules across the 10 TCR clonotypes.

**Supplemental Figure 6.**
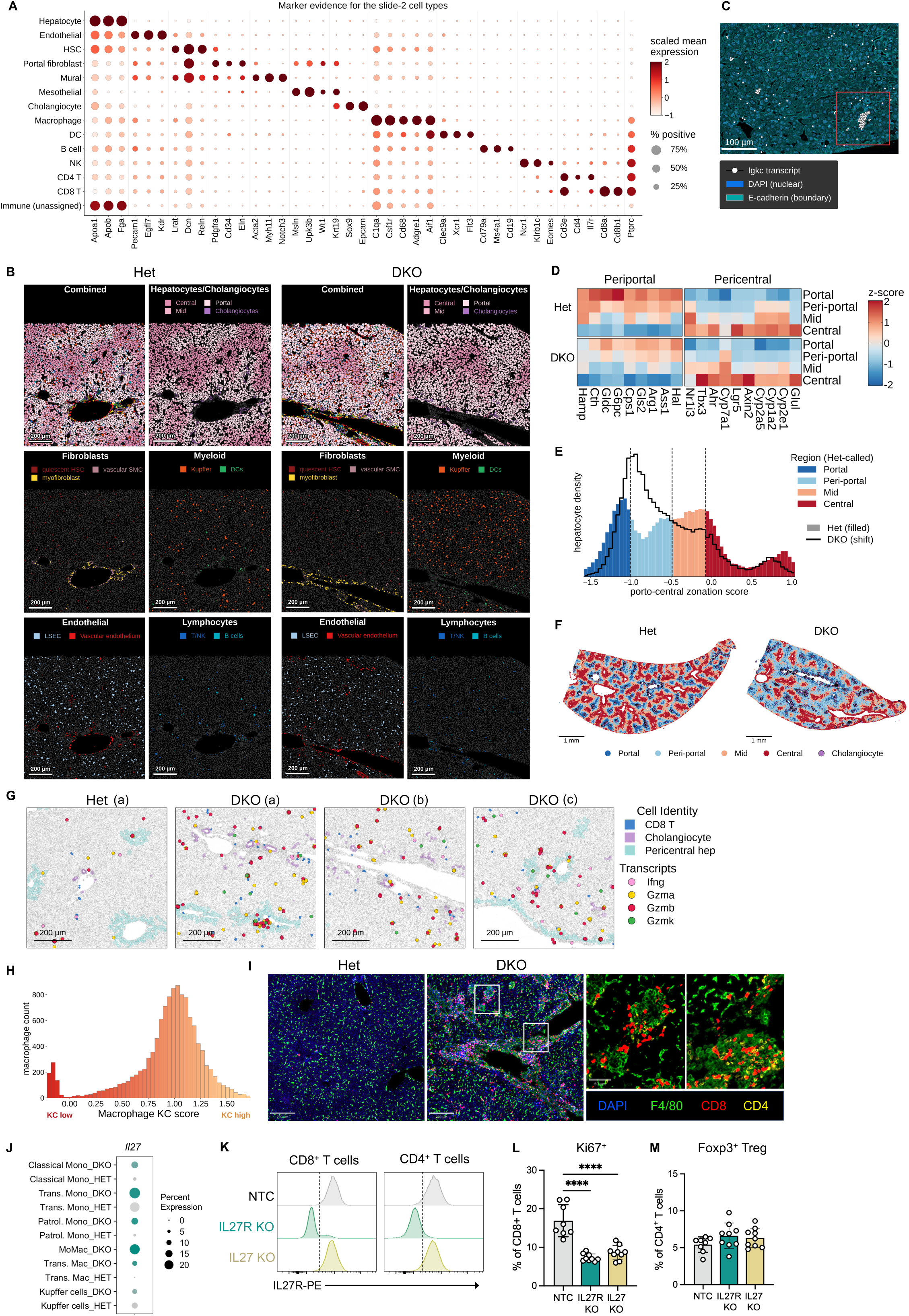
(A) Expression of genes that define clusters identified by Xenium spatial transcriptomics shown in figure 4A (B) Representative spatial transcriptomic image of DKO liver section showing transcript distribution of Igkc, marker gene of plasma cells. (C) Representative spatial transcriptomic images showing the distribution of the indicated parenchymal and non-parenchymal cell populations in Het and DKO liver sections. (D) Heatmap showing expression of genes defining periportal and pericentral hepatocytes. (E) Distribution of hepatocytes along the portal-to-central zonation axis in Het and DKO livers. Filled histograms show Het hepatocytes, and the black outline shows the distribution of DKO hepatocytes. Dashed lines indicate boundaries defining indicated regions based on the Het liver. (F) Spatial mapping of the zonation pattern in Het and DKO liver sections (Scale bars, 1 mm). (G) Representative spatial transcriptomic images showing the distribution of indicated T cell-associated transcripts in the same regions are same as shown in figure 4B. (H) Distribution of macrophages according to continuous Kupffer cell (KC) score calculated based on mean expression of *Cd5l* and *Vsig4*. (I) Representative immunofluorescence images (scale bar, 200um) of liver sections from Het and DKO mice showing macrophage (Green), CD8+ (red) and CD4+ (yellow) T cell clusters. (J) Expression of Il27 across monocyte and macrophage subsets from Het and DKO livers identified by scRNA-seq. (K) Representative histogram comparing surface IL27R expression on circulating CD8+ and CD4+ T cells in bone-marrow chimeras 9-week post transplantation. (L) Frequencies of Ki67^+^ CD8^+^ T cells in the livers of bone-marrow chimeras 9-week post transplantation (n = 8=9 per group). (M) Frequencies of Tregs in the livers of bone-marrow chimeras 9-week post transplantation. Data shown in (K) – (M) are pooled from at least 2 independent experiments and represented as mean ± SEM. *p < 0.05, **p < 0.01, ***p < 0.001, and ****p < 0.0001

**Supplemental Figure 7.**
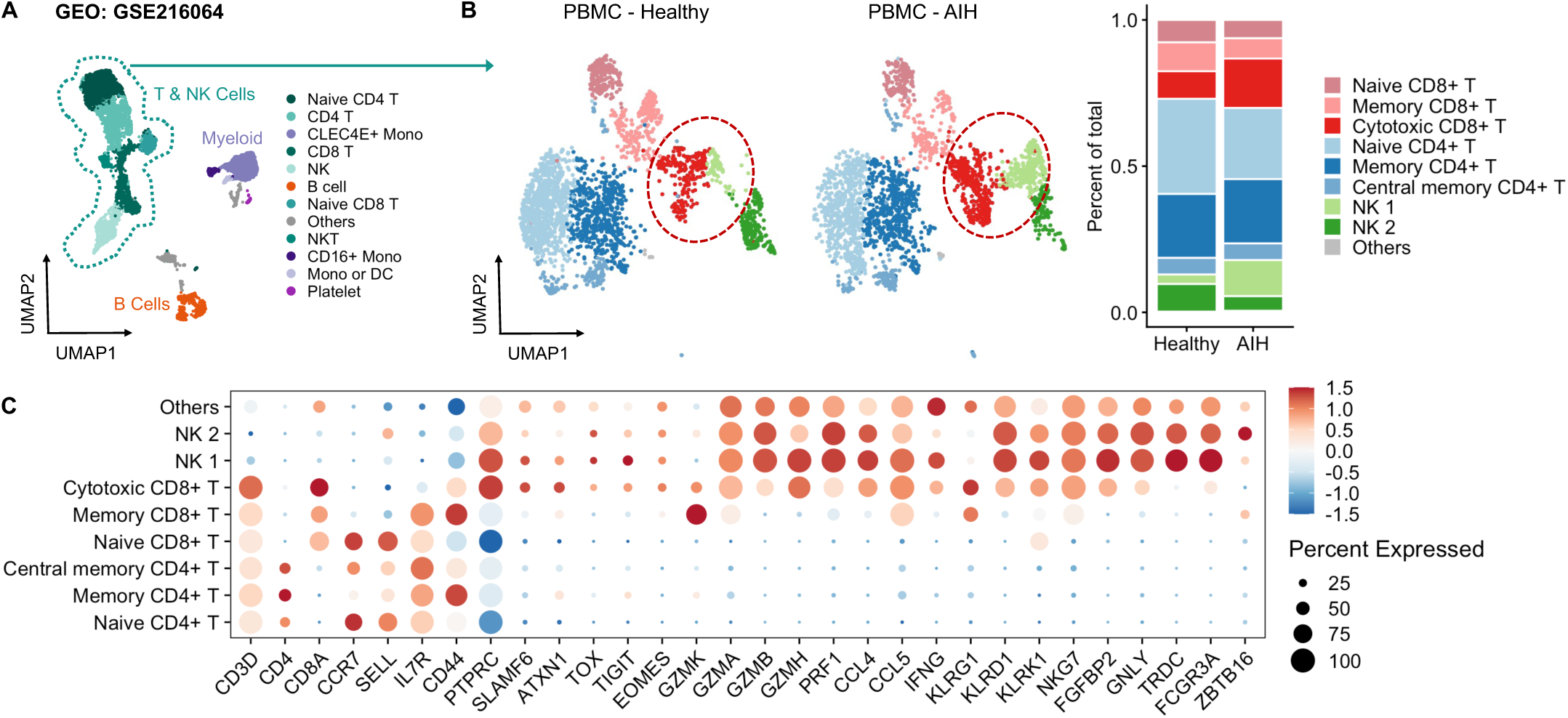
(A) UMAP plot of peripheral blood mononuclear cells (PBMCs) from healthy controls and patients with autoimmune hepatitis (AIH) in a published scRNA-seq dataset (GEO: GSE216064). (B) UMAP plots and relative abundance of T and NK cell subsets in healthy controls and patients with AIH. (C) Expression of cell-type-defining and cytotoxic effector genes across the T and NK cell subsets identified in (B).

